# Cell motility enhances metabolic coupling in spatially structured microbial communities

**DOI:** 10.64898/2026.04.29.720302

**Authors:** Miaoxiao Wang, Olga T. Schubert, Martin Ackermann

## Abstract

Metabolic interactions are central to the functioning of microbial communities. In spatially structured environments, such interactions typically require close physical proximity between partner cells. However, cell division drives spatial segregation of interaction partners, as the cells emerging from division remain adjacent and form clonal clusters, weakening these interactions. Here, we hypothesized that active cell motility can reduce this spatial segregation by enabling cells to leave clonal clusters and thereby enhance metabolic interactions. We tested this hypothesis by performing time-lapse single-cell imaging of a synthetic cross-feeding consortium growing in microfluidic chambers. We found that surface motility disrupted clonal clustering, enhanced spatial intermixing, and consequently increased the growth rates of individual cells and community productivity. Individual-based simulations further revealed that this effect is robust across motility modes and a wide range of ecological and physiological parameters. Together, our findings demonstrate that even non-directed, random motility, without chemotactic sensing, is sufficient to enhance metabolic interactions by separating cells from their clonal lineages and repositioning them in proximity to their metabolic partners, thereby acting as a key driver of community functioning in spatially structured microbial systems.

## Introduction

Microbial communities play pivotal roles across virtually all habitats on Earth. These communities typically consist of numerous species that can affect each other’s activity, growth, and survival through the exchange of metabolites^1–3^. Such metabolic interactions are ubiquitous in natural microbial communities^1,3,4^, shape their assembly and functioning^6–8^, and profoundly impact the physiology, ecology, and evolution of community members^1,5^. Understanding how different factors shape these interactions—and thereby influence community assembly and function—is therefore of fundamental importance^6^.

In nature, microbial communities are typically spatially structured rather than well-mixed^7^. In these settings, the distance between individual cells significantly influences the strength of their realized interactions^8–10^. Secreted metabolites are diluted as they diffuse across space and are rapidly taken up by neighboring cells, such that the strength of metabolic interactions between cells decreases rapidly with increasing spatial separation^11^. Indeed, our previous work demonstrated that individual cells only interact effectively with neighbours located within a short distance, typically in the range of 3-10 μm^10^. While a high degree of spatial mixing and short average distances between cells from interacting populations would thus enhance interaction strength^12,13^, several biological and physical processes decrease mixing and separate interacting populations. An important driver of this separation is clonal clustering, the spatial aggregation of clonally related cells resulting from growth and division. Clonal clustering arises when the cells emerging from successive division events remain close to each other, leading to spatial clusters of genetically identical cells. This process spatially separates cells potentially engaged in metabolic exchange and thereby weakens their realized interactions (Figure 1a).

**Figure 1.**
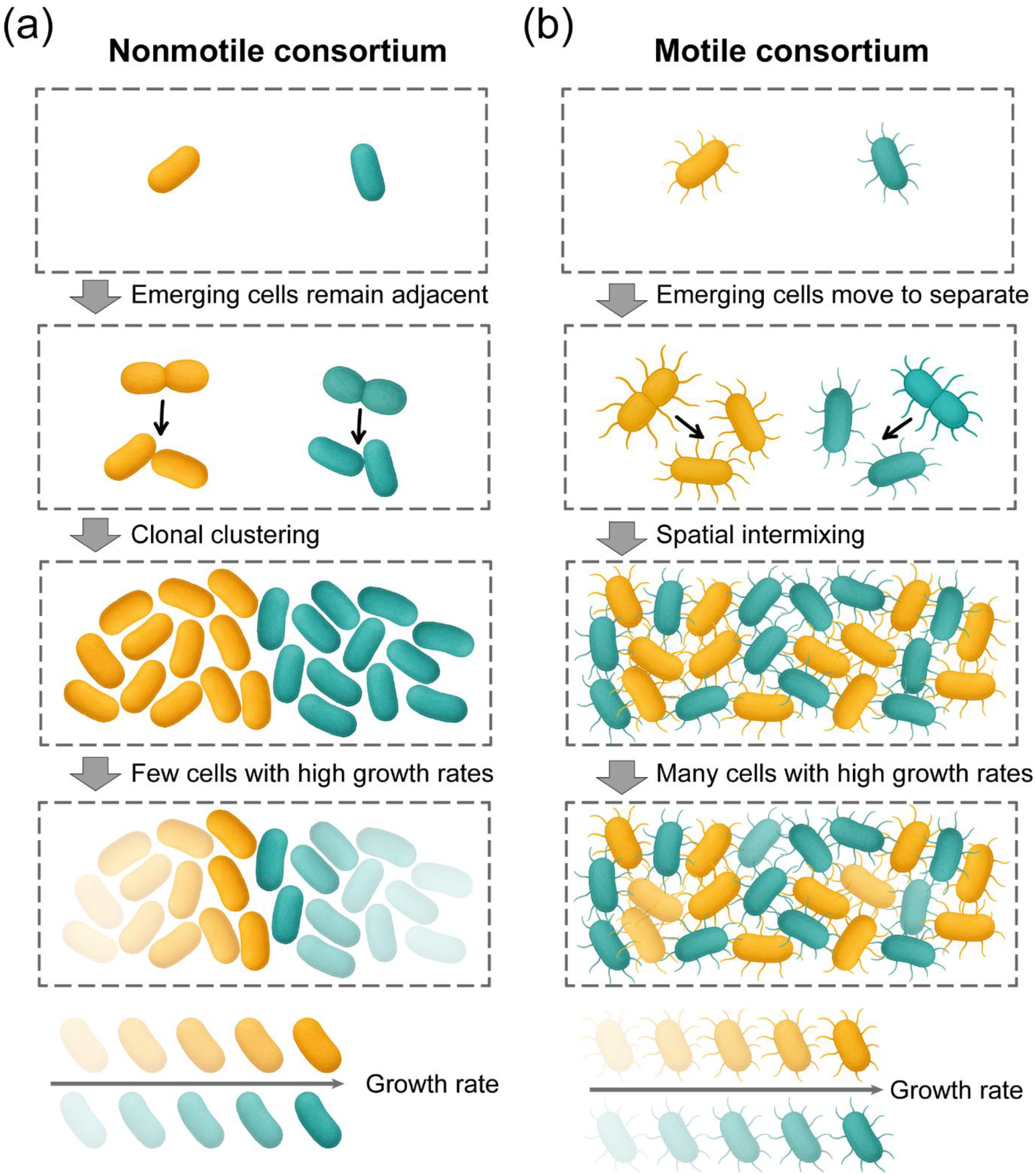
Conceptual overview of how motility affects the spatial arrangement of cells in a synthetic consortium consisting of two obligately cross-feeding genotypes. We worked with synthetic communities consisting of two genotypes that exchange essential metabolites within a spatially structured environment. (a) In the absence of motility, cells form clonal clusters, as cells remain close to each other after division. This clustering increases the average distance between genotypes and weakens metabolic coupling, resulting in decreasing growth rates for cells further away from the other genotype. (b) When cells are motile, the formation of clonal clusters is reduced because cells can move away from their clonal cluster. As a result, intermixing between genotypes and metabolic coupling increases, resulting in enhanced growth. The fourth row shows growth rates encoded by opacity, with more opaque colours indicating higher growth rates.

Here, we hypothesized that active movement of cells can reduce this spatial segregation and thereby enhance metabolic interactions: motile cells can leave clonal clusters and get closer to partner cells, thereby increasing access to essential metabolites and achieving faster growth (Figure 1b). Previous studies have shown that cell motility can increase spatial intermixing^14,15^ and alter inter-population interactions^16–18^ in macroscale colonies. At the micrometer scales at which individual cells grow and move, abiotic constraints, such as diffusion gradients^19,20^ and limited physical space^21–23^, and biotic factors, including single-cell motility modes^24,25^ and cell-to-cell variability in metabolism and growth^26^, can fundamentally alter how cells encounter and interact with one another. Our goal was thus to test directly, at the level of individual cells, whether motility would allow cells to access more metabolic partners and whether this would increase the growth of individual cells and of the community as a whole.

For this, we engineered synthetic bacterial consortia with different levels of motility where the members are engaged in metabolic interactions through the exchange of two amino acids, tryptophan and histidine. Amino acids are cellular building blocks essential for growth, and their cross-feeding is widespread in natural microbial communities^27–31^. Tryptophan and histidine are two amino acids frequently exchanged, for example, in anaerobic methanogenic consortia^32^ and the human gut microbiome^29^, potentially owing to their high biosynthetic costs^33^. Accordingly, synthetic consortia composed of tryptophan and histidine auxotrophs have been widely used to examine the ecological and evolutionary consequences of metabolic coupling between microbial partners^34–39^. With our engineered cross-feeding consortia of motile and non-motile auxotrophs, we performed time-resolved, single-cell imaging using microfluidic growth chambers to compare their growth dynamics. We then developed a mathematical model to identify the key factors that determine the extent to which cell motility enhances metabolic interactions.

## Results

### Cell motility enhances metabolic coupling in the synthetic consortia

To investigate whether cell motility enhances metabolic interactions, we constructed synthetic consortia of genetically engineered *Pseudomonas aeruginosa* (Figure 2a and b). To enforce reciprocal metabolic interactions between two genotypes, we deleted the *trpB* or *hisD* gene to generate tryptophan and histidine auxotrophs. As expected, neither strain grew alone in minimal medium without supplementation of amino acids (Methods), but both grew in co-culture, indicating that they engaged in mutualistic interaction by exchanging amino acids (Figure S1). To increase cross-feeding, we added the amino acid precursors indole (for tryptophan) and histidinol (for histidine) to the cultures, which leads to overproduction of the respective amino acids by the producer strain but does not rescue the auxotrophs (Figure S1).

**Figure 2.**
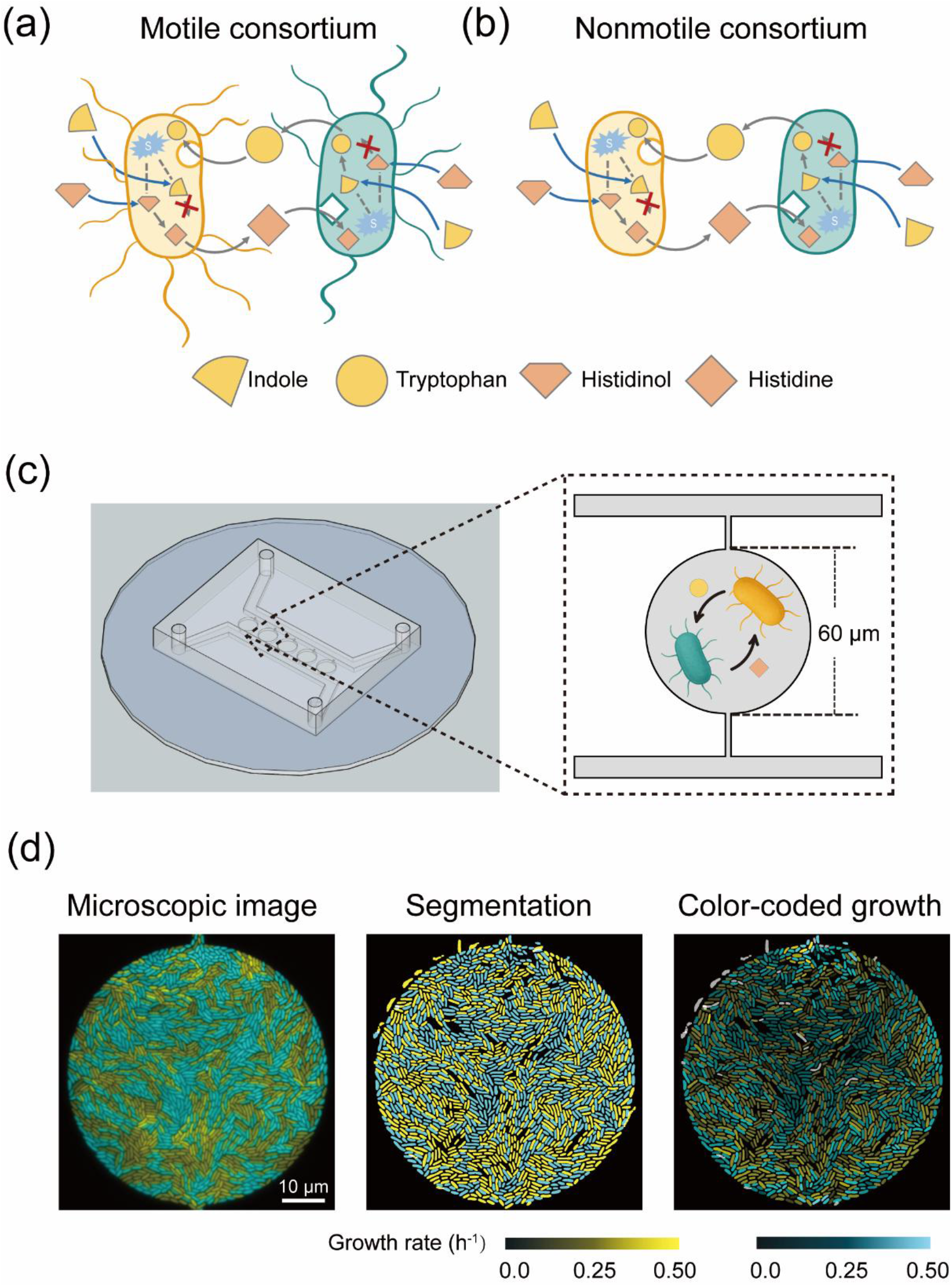
Design of synthetic consortia and microfluidics-based live-cell imaging experiments to investigate the effects of motility on metabolic interactions and growth. (a-b) Tryptophan and histidine auxotrophs of *Pseudomonas aeruginosa* PAO1 were constructed by deleting the biosynthetic genes *trpB* and *hisD*, respectively. Direct pairing of the two auxotrophs forms a motile consortium (a), while additional deletion of *pilA* and *filC* disabled twitching and swimming of the cells, yielding the nonmotile consortium (b). The amino acid precursors indole (for tryptophan) and histidinol (for histidine) were added to increase the respective amino acid production, thereby enhancing cross-feeding strength (Figure S1). (c) Consortia were cultured in monolayers in microfluidic chambers (0.5 μm high). Shown is a 60 μm circular chamber, which is connected to feeding channels by 3 μm passages that limit cell escape but allow medium in- and outflow. With this design, we could quantify the population growth rate by tracking the area occupied by cells over time (cells are not drawn to scale).(d) Strains were fluorescently labeled (yellow, cyan) for time-lapse imaging (left). Automated segmentation and tracking (middle) enabled the quantification of single-cell growth rates, which were visualized by color coding, with brighter hues indicating faster growth (right) and grey color indicating mis-tracked cells.

*Pseudomonas aeruginosa* exhibits two major forms of motility: type IV pili-mediated twitching on surfaces^40,41^ and flagella-driven swimming in liquid or near-surface^25,42^. To control motility in our experiments, we further engineered the auxotrophic strains to lack key motility genes. Specifically, we deleted *pilA*, which encodes the major pilin subunit essential for type IV pili and twitching motility^40,41^, and *fliC*, which encodes a flagellar hook–filament junction protein required for flagellum assembly and swimming^43^. These deletions disable the motility of these strains but do not affect their mutualistic growth in the well-mixed cultures (Figure S1). Using these mutant strains, we constructed two consortia: (1) a motile consortium consisting of the two motile auxotrophic strains (Figure 2a) and (2) a nonmotile consortium consisting of the two nonmotile auxotrophic strains (Figure 2b). We grew these consortia in monolayers in custom-designed microfluidic chambers, which are connected to feeding channels by 3 μm passages that limit cell escape but allow medium in- and outflow (Figure 2c). This setup allowed us to track single cells over time and use automated image analysis to quantify growth at both population and single-cell levels (Figure 2d; Methods).

Visual inspection of the images recorded over time indicated that the nonmotile consortia formed segregated spatial patterns, due to clonal clustering (Figure 3a; Figure S2; Supplementary Video 1-5). In contrast, cells in the motile consortium formed more intermixed patterns (Figure 3a; Supplementary Video 1-4). To quantify this difference in intermixing, we defined an intermixing index as the mean fraction of partner cells among all cells within a 5 µm neighborhood (Methods). Using this metric, we found that the motile consortium consistently exhibited higher intermixing levels than the nonmotile consortium throughout the time course (Figure 3b; Figure S3; Mann–Whitney U test: *p* < 0.01). This result indicates that cell motility disrupts clonal clustering and reduces the average distance between interacting populations.

**Figure 3.**
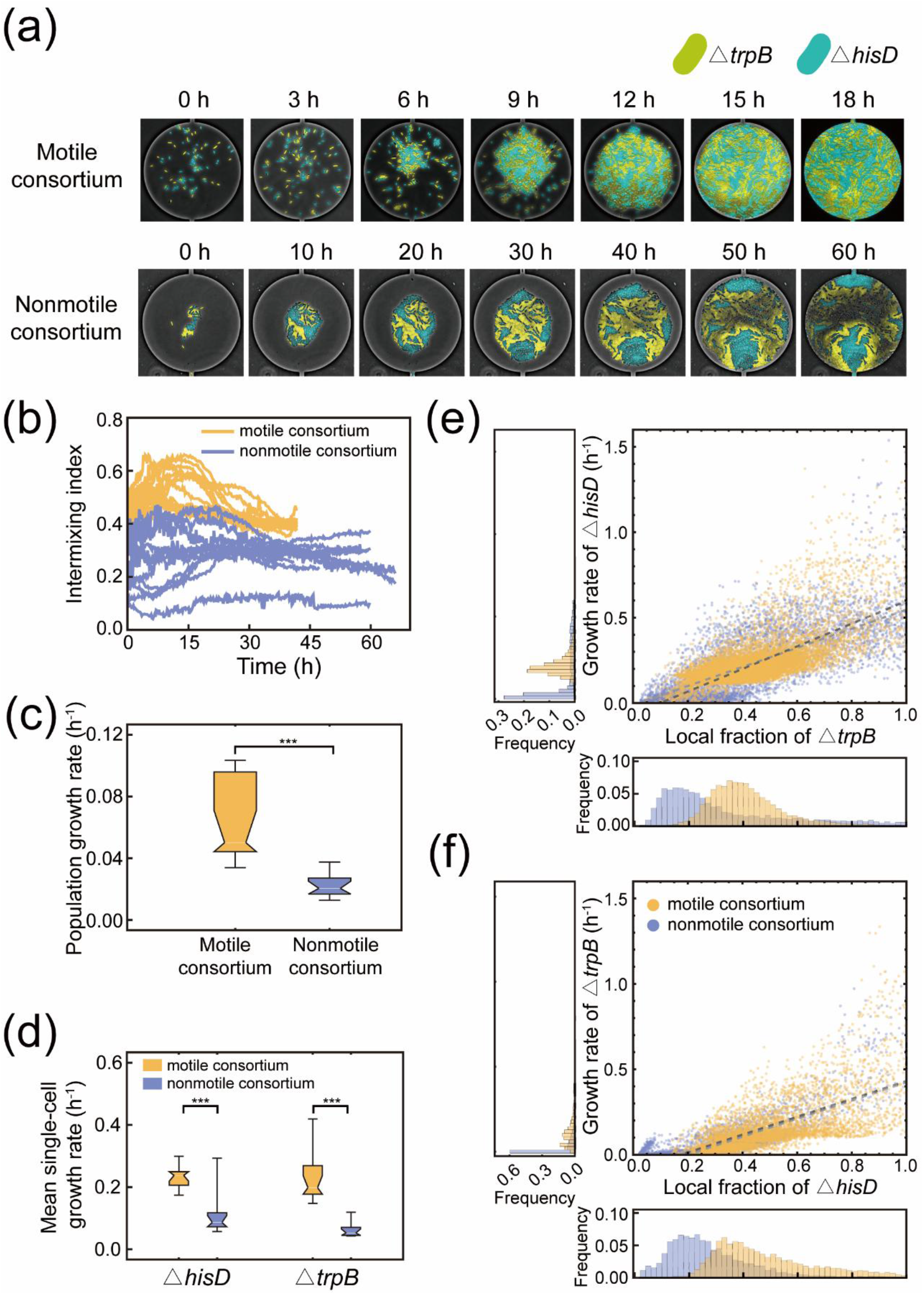
Live-cell imaging in microfluidic growth chambers shows that motility enhances metabolic interactions and growth in a consortium of obligately cross-feeding strains. (a) Representative images from time-lapse microscopy of motile (top) and nonmotile (bottom) consortia in the 60 μm circular microfluidic chambers. Nonmotile cells formed clonal clusters and segregated spatial patterns, whereas motile cells created more intermixed arrangements. Tryptophan auxotrophs are shown in yellow, while histidine auxotrophs are shown in cyan. (b–d) Comparison of intermixing index (b), population-level growth rates (c), and mean single-cell growth rates within each chamber (d). Data represent 14 chambers with motile consortia and 12 chambers with nonmotile consortia, pooled from three independent experiments. Statistical significance was assessed by Student’s t-test with individual chambers treated as independent replicates: *: *p* < 0.05; **: *p* < 0.01; ***: *p* < 0.001. Images were acquired at 12-min intervals. (e-f) Δ*hisD* cells (e) and Δ*trpB* cells (f) in both the motile and nonmotile consortia grew faster as the fraction of partner cells increased within the local neighborhood of a focal cell (5 μm radius). Yellow dots represent single cells in the motile consortia (16,104 Δ*hisD* cells and 5,304 Δ*trpB* cells), while blue dots represent single cells in the nonmotile consortia (7952 Δ*hisD* cells and 4,941 Δ*trpB* cells). The light grey line represents the linear regression of the motile cells with the coefficient of determination (R^2^ = 0.461 for Δ*hisD* cells in (e), and R^2^ = 0.438 for Δ*trpB* cells in (f)), while the dark grey line represents the linear regression of the nonmotile cells with the coefficient of determination (R^2^ = 0.650 for Δ*hisD* cells in (e), and R^2^ = 0.641 for Δ*trpB* cells in (f)). The histograms denote the frequency distributions of the local partner frequency and the single-cell growth rates. The same analyses for other chamber geometries are shown in Figure S3-S6.

To investigate whether this reduction in interpopulation distance enhances the growth of the motile consortium, we quantified population-level growth rates (Figure S4). Our analysis indicated that the motile consortium grew significantly faster than the nonmotile consortium (Figure 3c; Student’s t-test: *p* < 0.01). The motile consortium reached a maximum specific growth rate of 0.066 ± 0.029 h^−1^ (mean ± standard error of the mean, SE), compared with 0.031 ± 0.014 h^−1^ (mean ± SE) for the nonmotile consortium. We also quantified single-cell growth rates in both consortia (Methods). Because rapid movement of cells during early stages precluded reliable tracking, we compared single-cell growth rates only after ∼65% of the chamber space was occupied (corresponding to ∼24 to 30 hours after inoculation), when the movement speed of motile cells decreased due to spatial constraints (Supplementary Video 6). At this stage, cells in the motile consortium exhibited significantly higher average growth rates than those in the nonmotile consortium (Figure 3d; Student’s t-test: *p* < 0.05), consistent with our analysis at the population level. To test the robustness of this effect, we repeated the experiments in microfluidic chambers with three other geometries and found that the growth advantage of the motile consortium was consistent across all of them (Figure S3; Student’s t-test: *p* < 0.01). Overall, our findings suggest that cell motility strongly enhances overall biomass productivity in our synthetic consortia.

We next assessed whether the higher growth rates of motile cells can be explained by increased spatial mixing, i.e., an increased fraction of partner cells within the local neighborhood of a focal cell (5 µm radius). If this were the case, we would expect that both motile and nonmotile consortia show the same relationship between growth rate and local partner fraction. However, motile cells would experience higher local partner fractions and therefore more of them would have higher growth rates. Consistent with this expectation, across replicates and different chamber geometries, we observed exactly that: Cells in the motile consortium generally experienced higher local partner fraction and higher growth rates than those in the nonmotile consortium (Figure 3e-f; Figure S5; Student’s t-test: *p* < 0.01), and the association between single-cell growth rate and local partner fraction was highly similar in both consortia (Figure 3e-f; Figure S5). While the statistical analysis shows that the fitted linear regressions are not identical between motile and nonmotile cells (Chow test ref: *p* < 0.001), null model analyses indicate strong similarity: for Δ*hisD* cells, similarity exceeds that of ∼91.9 % of null realizations, and for Δ*trpB* cells, it exceeds that of ∼92.0 % of null realizations (Figure S6; Table S1). Together, these analyses indicate that increased spatial intermixing driven by motility explains most of the growth advantage of motile cells, although additional motility-related effects likely play a secondary role.

### Mathematical modeling identifies key factors determining the positive effects of cell motility

Our experiments establish that motility can strengthen metabolic interactions by disrupting clonal clustering, reducing the physical separation between interacting populations and thereby increasing the growth of individuals and the productivity of the consortium. However, these experiments were conducted under specific environmental conditions and culture settings, leaving open important questions about the generality and underlying mechanisms of this effect. In particular, it is unclear whether the benefits of motility depend on particular motility modes, the specifics of metabolic interactions, or growth kinetics. To address these aspects, we developed an individual-based model that allows for systematic exploration of the parameter space and mechanisms governing how cell motility enhances metabolic interactions.

#### Design and evaluation of the individual-based model

Our individual-based model was built to address two questions: (1) Does cell motility increase metabolic interactions only under specific conditions or more generally? (2) What are the key parameters that determine the magnitude of this effect? To describe the metabolic exchange, growth, and cell division, we adapt the modeling framework of our previous works^10,13^, which places cells on a 40 × 40 lattice with up to one cell per lattice position and is based on the following assumptions: (1) cells actively take up amino acids; (2) cells passively leak amino acids; (3) amino acid consumption is proportional to cell growth rate; (4) cells produce amino acids to maintain a constant internal concentration through regulation of production rates; (5) cell growth is limited by the amino acid they cannot synthesize, following Monod kinetics; and amino acids diffuse in the extracellular environment with a rate proportional to the concentration gradient of the amino acid; (7) Cells divide when their biomass reaches a threshold, after which one cell remains in the original grid, while the other cell moves to one directly adjacent lattice position. If this position is already occupied, one of the two cells on this position is selected at random and removed. Within this framework, four parameters are directly related to growth and metabolic interactions: *D*, the diffusion coefficient of the amino acids; *rL*, the leakage rate; *rU*, the uptake rate; and *gE*, the intrinsic growth rate of the cell. Mathematical details are provided in the Supplementary Information.

To analyse the effects of motility on spatial mixing and metabolic interactions, we implemented the following three motility modes into the model (see Supplementary Information for details). The first mode is a nonmotile state where cells remain in a given position. The second mode is a random motility pattern, characterized by three parameters: movement probability (*Mp*), reorientation noise (*Rn*), and movement speed (*Ms*). The third mode is an aggregative motility pattern based on our experimental observations, where we saw that cells move rapidly during the early stages of colonization, then start aggregating toward the center and gradually become sessile (Figure 4a; Supplementary Video 7). This pattern is consistent with signal-guided twitching motility in *P. aeruginosa*^44,45^, where cells constitutively secrete a quorum-sensing exopolysaccharide signal and stop moving once the local concentration of this quorum-sensing signal exceeds a threshold (denoted by *St* in our model). We implemented this mechanism into our model to test whether this would increase the fit to our experimental observations.

**Figure 4.**
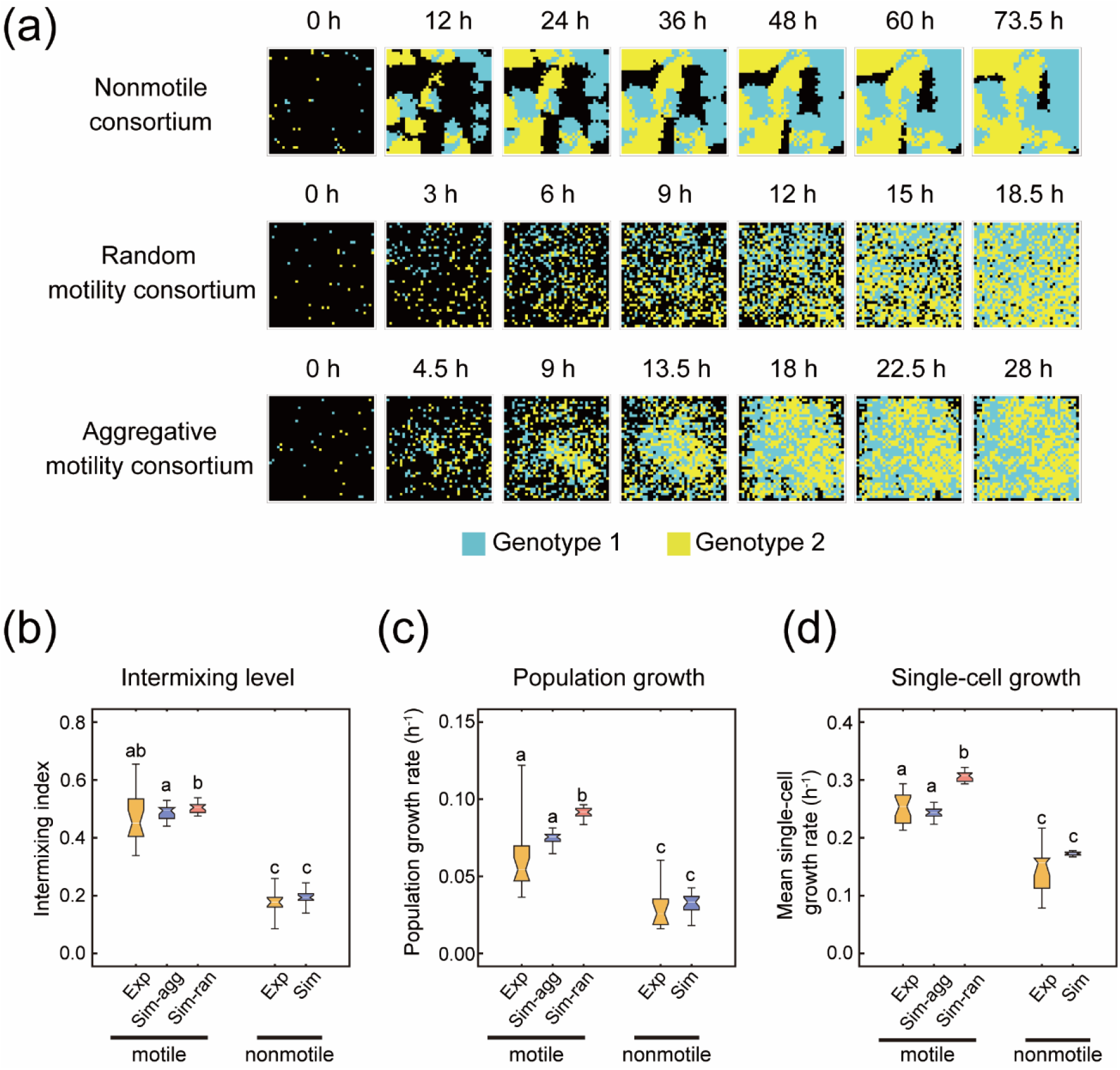
Individual-based simulations reproduce experimental observations. (a) Representative dynamics of nonmotile (top), random motility (middle), and aggregative motility (bottom) consortia simulated with default parameter values shown in Table S2. (b–d) Comparison of intermixing index (b), population-level growth rates (c), and mean single-cell growth rates (d) between simulations and experiments in 40 μm square chambers. Simulations represent 20 replicates for each motility scenario; experimental data represents 21 and 14 chambers with motile and nonmotile consortia, respectively, that were recorded in three independent experiments. ‘Exp’ denotes the data derived from our microfluidic experiments, while ‘Sim’ denotes the simulation data. ‘agg’ denotes the aggregative motility mode in the simulation, while ‘ran’ denotes the random motility mode. In the notched box-and-whisker plots, the white center line indicates the median; the notch around the median represents the 95% confidence interval of the median; the lower and upper bounds of the box denote the first (Q1, 25%) and third (Q3, 75%) quartiles, respectively; and the whiskers extend to the most extreme values within 1.5 × IQR. Statistical significance was assessed by Student’s t-test. Different lowercase letters (a, b, c) indicate statistically significant differences between groups (*p* < 0.05).

To evaluate whether the model reproduces the patterns observed in our microfluidic live-cell imaging experiments, we ran it using experimentally derived parameters (Table S2). The simulations with nonmotile strains reproduce the growth and clonal cluster formation observed in the experiments using the nonmotile synthetic consortium (Figure 4a; Supplementary Video 7). Simulations incorporating the random motility mode do not fully reproduce the experimental observations, yielding higher levels of spatial intermixing, as well as faster population and single-cell growth than those experimentally observed in the motile consortia (Figure 4b-d). In contrast, simulations based on the aggregative motility mode recapitulate the experimental dynamics: cells move rapidly during early stages, then aggregate toward the center and gradually become non-motile (Figure 4a; Supplementary Video 7). Quantitatively, these simulations most closely match the experimental results in the 40 μm square microfluidic chambers (Figure 4b-d; Supplementary Video 4; Supplementary Video 7).

#### Cell motility enhances metabolic interactions across different conditions

To examine whether cell motility enhances metabolic interactions across different conditions, we varied the parameters describing the initial density, growth, and metabolic interactions (Figure 5a). We generated 300 parameter sets spanning a wide range of values for *D, rL, rU*, and *gE* based on empirical estimates reported in previous studies (Table S2; Supplementary Information). For each set, we ran simulations under five initial cell densities and eight initial spatial distributions, resulting in 12,000 simulations for each motility pattern.

**Figure 5.**
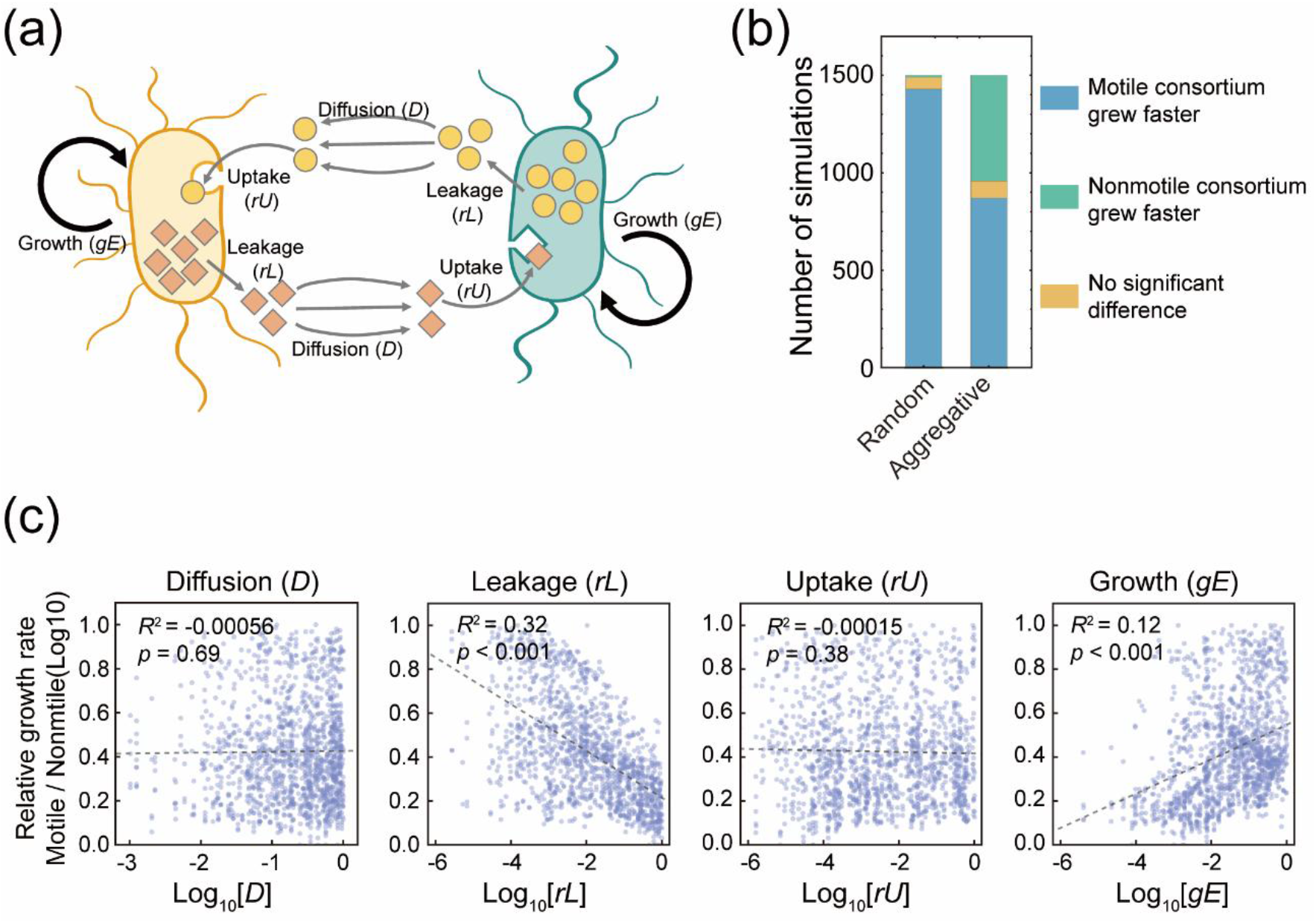
Effects of intrinsic growth rate and metabolic interaction parameters on the positive effects of cell motility. (a) Schematic representation of the model assumptions capturing the metabolic exchange between two auxotrophic mutants. A cell with a particular amino acid auxotrophy synthesizes an internal pool of another amino acid, which leaks into the environment at a rate *rL*, diffuses across the extracellular habitat at a rate *D*, and is taken up by a cell with an auxotrophy for this other amino acid at a rate *rU*, and vice versa. Both auxotrophic mutants grow following Monod kinetics with a maximum growth rate of *gE*. (b) Summary of whether motile consortia exhibited higher population growth rates than nonmotile consortia for both types of motility. (c) Effects of *D, rL, rU*, and *gE* on the relative population growth rates of the random motility consortium compared with the nonmotile consortium.

Consistent with experimental observations, motility increased the intermixing level of the consortium in most simulations. Specifically, random motility increases the intermixing level in 80.2% (9,632/12,000) of the simulations, and aggregative motility increases the intermixing in 63.6% (7,637/12,000) (Figure S7a). Higher intermixing levels led to more uniform spatial distributions of metabolites (Figure S8a–b) and increased the average single-cell growth rates (Figure S7b; Figure S8c–d). Consistent with these effects at the single-cell level, the consortium’s overall productivity was also increased: The random motility consortium exhibited higher population-level growth rates in 97% of simulations (11,658/12,000), whereas the aggregative motility consortium grew faster in 60.8% (7,297/12,000; Figure 5b; Figure S7c; Figure S8e–f). Together, these results demonstrate that the mechanism by which cell motility enhances metabolic interactions and growth (through the promotion of spatial intermixing) operates robustly across a wide range of conditions, even though the aggregative motility pattern reduces this effect.

#### Leakage rate and intrinsic growth rate determine when motility enhances metabolic interactions

Having established that cell motility can enhance metabolic interactions, we next asked under what conditions this effect emerges and how ecological (e.g., initial cell density) and physiological (e.g., uptake and leakage rates of exchanged metabolites, as well as intrinsic growth rate) factors affect its magnitude. These factors could, in principle, modulate the extent to which motility-driven intermixing translates into increased growth of the consortium.

To address these questions, we analyzed our simulation results derived from the above 300 parameter sets. A correlation analysis indicated that the positive effect of motility on growth was more pronounced when cells leaked metabolites more slowly and possessed higher intrinsic growth rates (Figure 5c; Figure S9). First, slow leakage rates restricted the spatial range over which cells interact^10,46,47^, thereby increasing the effect of intermixing through motility. Second, motility can only increase intermixing once cells start encountering other cells, whereas intrinsic growth rate controls how fast the system exits this low-density regime (Figure S10a). At early stages, cell density is low, and most cells are surrounded by empty space, so motility rarely leads to encounters with other cells and has little effect on increasing spatial mixing. As cell density increases, cells begin to encounter nearby neighbours, and motility can then promote the encounter of interacting partners. A higher intrinsic growth rate has two effects. First, it shortens this low-density phase and allows motility to become effective earlier in promoting partner encounters. Consistent with this interpretation, the relative time point *Tm*, at which motile consortia begin to experience more partner encounters than nonmotile consortia, decreased with increasing *gE* (Figure S10c-d). Second, once such encounters occur, cells with higher intrinsic growth rates can more rapidly translate improved access to partner-derived amino acids into biomass. In other words, differences in local partner availability caused by motility are converted more quickly into differences in realized growth. Together, these results show that the extent to which motility enhances growth depends on physiological parameters that shape both the spatial range of interactions and how efficiently cells convert partner-derived metabolites into biomass.

#### Positive effects of motility remain robust to different motility parameters under random motility, but not aggregative motility

Beyond ecological and physiological factors, differences in motility modes and in how cells move (described here by kinematic parameters) can also modulate how motility influences metabolic interactions by altering spatial intermixing. To quantify these effects, we systematically explored parameter space by generating 50 additional combinations of kinematic parameters—movement probability (*Mp*), reorientation noise (*Rn*), and movement speed (*Ms*; Figure 6a). Each combination was tested using 20 random parameter sets spanning different values of ecological and physiological factors. Using these values, we ran 1,000 simulations for each of the random and aggregative motility patterns, and 20 simulations of the nonmotile scenario as a control.

**Figure 6.**
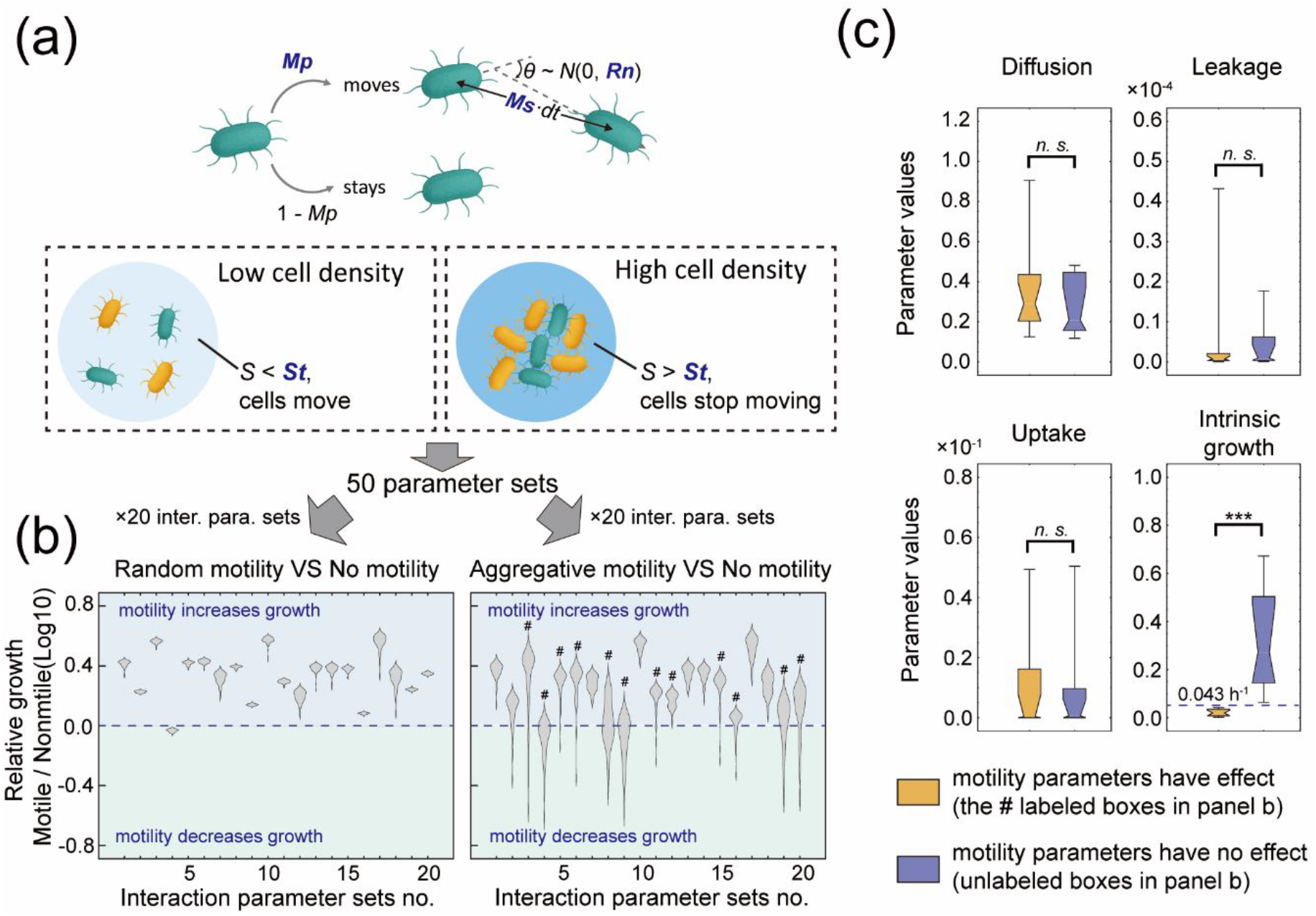
Effects of motility-related parameters on the positive effects of cell motility on growth. (a) Schematic representation of the motility assumptions in the model. At each time step, a cell moves with probability *Mp*. If cell movement occurs, cell orientation is updated by a turning angle *θ* drawn from a normal distribution N (0, *Rn*), where larger *Rn* values generate more randomized, diffusive trajectories. The moving distance is drawn from a normal distribution with an expectation of *Ms·dt*, where *Ms* is the movement speed, and *dt* is the time-step. To simulate aggregative motility, we assumed that all cells continuously secrete a diffusible QS signal that accumulates in the extracellular environment. At each simulation step, every cell senses the local concentration of this signal within its grid. When the sensed concentration exceeds a predefined threshold (*St*), the cell ceases movement for that iteration. (b) Relative population growth rates of motile versus nonmotile consortia across 20 growth- and interaction-related parameter sets; each violin shape represents results from 50 motility-related parameter combinations. For the aggregative motility scenario, thirteen parameter sets in which motility-related parameters altered whether motility increases community growth are marked with #. (c) Comparison of growth- and interaction-related parameter values between the thirteen sensitive sets and the seven insensitive sets determined in the right panel of (b). The intrinsic growth rate threshold (*gE*_*thres*_ *=* 0.043) distinguishing the two cases is shown in the lower right panel.

When cells exhibited random motility, our analysis indicates that kinematic parameters rarely determine whether motility enhances metabolic interactions (Figure 6b). They only affect the extent to which motility confers a positive effect. In contrast, the effect of aggregative motility on community growth is more sensitive to changes in kinematic parameters (Figure 6b). Our analysis reveals that the effect of kinematic parameters on aggregative motility depends on the intrinsic growth rate (*gE*): above a threshold (0.043 h^-1^ in Figure 6c), varying *Mp, Rn*, and *Ms* does not alter whether aggregative motility enhances community growth, whereas below this threshold it does (Figure 6c). This growth rate threshold decreases with the quorum-sensing threshold, *St* (Figure S11a), as faster growth leads to faster accumulation of the quorum-sensing signal, inducing cessation of movement (Figure S11b) and the formation of aggregates (Figure S11c). This early arrest of movement limits the ability of motility to increase spatial intermixing (Figure S11d). In contrast, slower intrinsic growth delays signal accumulation and prolongs the motile phase (Figure S11d). As a result, the formation of aggregation becomes more sensitive to differences in kinematic behaviours of the cells (Figures S11e–f), which significantly influences community growth (Figure S11g). For example, specific kinematic parameter conditions increase the probability of cell encounter, leading to the formation of a single aggregate^48^, which reduces the level of intermixing (Supplementary Video 8-10). Overall, these results show that random motility robustly enhances metabolic coupling across diverse conditions, whereas aggregative motility can either enhance or reduce metabolic coupling depending on intrinsic growth rate and quorum-sensing thresholds.

## Discussion

In this study, we combined microfluidics-based quantitative single-cell analysis with individual-based modeling to examine how cell motility affects metabolic interactions in microbial communities. Our experiments provide quantitative evidence that cell motility enhances metabolic interactions by disrupting clonal clustering and reducing the average intercellular distance between interacting populations. The individual-based model allowed us to probe the generality of this mechanism beyond the specific experimental conditions and to identify the determinants of whether motility enhances metabolic coupling. The simulations revealed that the positive effects of motility are primarily shaped by several key factors, including initial cell density, leakage rate, and intrinsic growth rate.

Recent studies highlight the ecological importance of cell motility in shaping microbial interactions as well as the assembly and function of microbial communities, for example, during the colonization of marine particles^49–52^, the phycosphere^53–55^, soil particles^56,57^, the infant gut^58^, and cheese rind^59^. Many of these studies emphasize that motility enables microorganisms to explore nutrient-rich niches, with motile microorganisms often dominating during the early phases of colonization. Our work highlights another role of motility in community ecology, namely that it allows cells to leave clonal clusters and move into neighborhoods with more metabolic partner cells. This increases access to metabolites from these partners, increasing and balancing growth across individual cells, thereby improving community growth.

This mechanism also offers an explanation for the formation of microscale multispecies aggregates with high levels of intermixing often observed in natural habitats^60–64^, despite the counteracting effect of clonal clustering. For example, anaerobic archaea have been reported to form intermixed aggregates with their syntrophic bacterial partners^62,63,65,66^. Several studies found that genes encoding motility-associated appendages such as flagella, archaella, and pili are strongly upregulated during the development of such aggregates^65–68^, suggesting an important role of motility in establishing these intermixed structures. More generally, our results suggest that, in surface-associated systems, motility is shaped by the dominant biotic interactions. Motility is expected to be upregulated when cells depend on metabolic partners and reduced when growth in clonal groups is advantageous or when interactions with other species are antagonistic^69^. These patterns may arise through regulatory responses at the level of gene expression or through evolutionary adaptation to recurrent ecological conditions.

Our model reveals that the effect of motility on metabolic coupling depends on the specific mode of motility. Random motility enhanced metabolic coupling robustly across diverse ecological and physiological conditions, whereas aggregative motility increased coupling only when sufficient intermixing is achieved before aggregation. This distinction is especially important as many microbial species (e.g., *P. aeruginosa*^44,45^, *Bacillus subtilis*^70,71^, *Vibrio cholerae*^72^, *Myxococcus xanthus*^73^, and *Escherichia coli*^74^) were reported to display aggregative behaviors during the early stages of biofilm formation^75^. Our results further suggest that the timing of aggregation relative to cell growth is a key determinant of whether aggregative motility enhances metabolic interactions. In particular, delaying the onset of aggregation, for example, by slowing the accumulation of quorum-sensing signals that stops movement, can extend the motile phase and allow sufficient intermixing to occur before cells form aggregation.

Our study shows that chemotaxis is not required for motility to enhance metabolic interactions. Even in the absence of chemotactic sensing, motility can enhance metabolic interactions by disrupting clonal clustering and relocating cells into neighborhoods with metabolic partners. This finding is notable because chemotaxis is often considered necessary for cells to reach metabolic partners and access their metabolites ^53,54,76^. Indeed, numerous studies have shown that bacteria exhibit strong chemotaxis toward a wide range of metabolites that can serve as nutrients or biosynthetic building blocks. These metabolites include not only amino acids^77–79^ but also carbohydrates, organic acids, peptides, benzenoids, lipids, and even nucleotide precursors^78,80^. Several studies showed that metabolic exchanges between bacteria and phytoplankton can be stimulated by chemotaxis^53,80^. However, other studies suggest that chemotaxis is not universally beneficial. For example, one recent study reported that cross-feeding between bacterial and yeast auxotrophs was enhanced by the non-directional motility of bacteria but not further improved by chemotaxis^81^. These findings suggest that while chemotaxis may strengthen metabolic interactions in some contexts, it is not a prerequisite for motility to be advantageous. Its role in dense, spatially structured communities remains context-dependent and warrants further investigation.

In summary, our study provides a general explanation for why motility can enhance growth even in the absence of chemotaxis. We established a quantitative framework linking motility, spatial structure, and metabolic coupling in microbial communities. This framework will be valuable for guiding efforts to understand, predict, and engineer the dynamics and functions of microbial communities in both natural and synthetic ecosystems.

## Materials and Methods

### Construction of the bacterial strains

All bacterial strains used in this study were derived from the *pelA* mutant of *Pseudomonas aeruginosa* PAO1. The absence of *pelA* disrupts the production of the glucose-rich Pel exopolysaccharide^82^, thereby preventing rapid three-dimensional biofilm formation within the microfluidic chambers, which otherwise leads to chip clogging and failure of single-cell tracking. The *pelA* mutant served as the parental strain for constructing the auxotrophic strains used in the motile consortium, whereas a triple mutant lacking *pelA, filC*, and *pilA* was used to generate the strains for the nonmotile consortium. These mutant backgrounds were kindly provided by Dr. Luyan Z Ma (Institute of Microbiology, Chinese Academy of Sciences). Auxotrophic mutants were constructed by deleting either the *hisD* or *trpB* gene via a standard two-step allelic exchange method based on homologous recombination^83^. For fluorescence labeling, *mcherry* or *gfp* genes were inserted at a neutral chromosomal site using the mini-Tn7 transposon-based tagging system^84,85^. All of these strains were freely available upon request.

### Media and batch culture assays

Prior to experiments, the fluorescent-labelled strains were streaked on LB agar plates supplemented with 30 μg mL^−1^ gentamicin. A single colony was inoculated into 2 mL of MOPS medium^86,87^ containing 20 mM succinate as the sole carbon source and 150 μM of the corresponding amino acid required by the auxotroph. Cultures were grown for 8 h at 37 °C with shaking. For inoculum preparation, cells from the pre-cultures were washed twice with carbon-free MOPS medium and resuspended to an optical density (OD_600_) of 5.0. For the inoculum of the synthetic consortia, equal volumes of the two partner auxotrophic strains were mixed at a 1:1 ratio. Batch growth kinetics were assessed in 96-well microplates using a microplate reader (Biotek). 20 μL of inoculum was added to 980 μL of MOPS medium supplemented with 20 mM succinate, with or without the amino acid precursors histidinol (2.5 μg·mL^-1^) and/or indole (10 μg·mL^-1^), resulting in an initial OD_600_ of approximately 0.1. Because the deleted genes (*hisD* or *trpB*) catalyze the final conversion of the precursors histidinol or indole into histidine or tryptophan, respectively, supplementing these precursors does not rescue the growth of the auxotrophs in monoculture (Figure S1). However, in coculture, the added precursors enable the partner strain (which retains the intact biosynthetic gene) to overproduce the amino acid, thereby increasing the amount of histidine or tryptophan leaked into the environment and thus enhance cross-feeding efficiency. This suspension was distributed into six wells with 150 μL for each well for six replicates. Cultures were incubated at 37 °C for 96 h, and OD_600_ was recorded every 10 min.

### Microfluidics and time-lapse microscopy

Microfluidic experiments were performed as previously described^10,88^, with modifications in the chamber design and inoculation procedure. Cells were cultured and imaged in monolayer microfluidic chambers (0.5 μm in height) of four geometries: circular chambers with diameters of 40 μm or 60 μm, and square chambers with side lengths of 40 μm or 60 μm. Each chamber was connected to two feeding channels via 3-μm connecting passages that prevented cell escape while allowing nutrient diffusion (Figure S1). The feeding channels (22 μm in height, 100 μm in width) were continuously supplied with MOPS medium containing 20 mM succinate,2.5 μg·mL^-1^ histidinol, and 10 μg·mL^-1^ indole. Inocula of the synthetic consortia were prepared following the same procedure described above. To load cells into the chambers, the inlet of the upper feeding channel was sealed using a short tubing connected to a 1-mL syringe containing 500 μL of flow medium. Another syringe containing 500 μL of the inoculum was connected to the outlet port of the same channel. Gentle pressure was applied to push cells into the upper feeding channel. Because the inlet of the upper feeding channel was blocked, the cells were were forced to flow through the centre chambers to enter below feeding channel. Once a sufficient number of cells entered the chambers, the tubing setup was replaced with that described previously^10,88^, in which two syringe pumps (NE-300, NewEra Pump Systems) equipped with 50-mL syringes delivered fresh medium into the feeding channels at equal flow rates of 0.1 mL h^−1^. This flow ensured sufficient nutrient diffusion into the chambers while preventing accumulation or exchange of amino acids through the main channels. No growth was observed in chambers inoculated with only one auxotrophic strain. Time-lapse imaging was conducted using Olympus IX81 or IX83 inverted microscope systems (Olympus, Japan) equipped with automated stage controllers (Marzhauser Wetzlar, Germany), shutters, and laser-based autofocus systems (Olympus ZDC 1 and ZDC 2). Phase-contrast and fluorescence (EGFP and mCherry) images were captured every 12 minutes from multiple chambers in parallel on the same PDMS chip. Temperature during imaging was maintained at 37 °C using a cellVivo incubation system (Pecon GmbH, Germany) or a Cube incubation system (Life Imaging Services, Switzerland).

### Automated image analysis

Automated image processing was performed using the custom-developed software MIDAP (Microbial Image Data Analysis Pipeline; https://github.com/Microbial-Systems-Ecology/midap; version 0.3.18) and custom scripts written in Wolfram Mathematica (Version 14.0), available online (Github: https://github.com/WMXgg/Cell-motility-project-codes/tree/main/PA%20project%20Code/Expcodes). The TIFF image sequences were processed in MIDAP for cell segmentation, lineage reconstruction, and continuous cell tracking, following the workflow detailed in our previous study^89^. MIDAP produced both segmented image stacks and comprehensive tracking files containing metadata for each cell. Because the rapid cell motility in the motile consortium prevented reliable cell tracking during early growth, only segmentation analyses were first performed across the entire image series. These segmented images were then used to quantify (i) the total area occupied by cells and (ii) the spatial distribution of cells within chambers. The temporal change in the fraction of chamber area covered by cells was then fitted to a Gompertz growth model to estimate population-level growth rates. Cell spatial positions were used to calculate the intermixing index, which quantifies the degree of local spatial mixing between the two cell types. Specifically, for each Δ*hisD* cell, we counted the number of neighboring Δ*trpB* and Δ*hisD* cells within a fixed 5 μm radius and calculated the local fraction of Δ*trpB* cells against the total cells. The intermixing index at each time point was obtained as the mean of these ratios across all Δ*hisD* cells. These analyses were performed using the custom Wolfram Mathematica scripts. After approximately 65% of the chamber area became occupied, when cell motility had largely reduced, segmentation and tracking analyses in MIDAP were combined to extract single-cell growth information. For this analysis, 30 - 60 frames were analyzed for the motile consortium and 80 - 150 frames for the nonmotile consortium. The tracking files included cell IDs, centroid coordinates, cell areas, major axis lengths, and fluorescence intensities for both channels, enabling quantitative single-cell analyses. Single-cell growth rates were estimated by fitting the time-dependent changes in major axis length (used as a proxy for cell size) as previously described^75^, using a custom Wolfram Mathematica script.

### Construction, simulation, and analysis of the individual-based model

The mathematical framework was adapted from our previously developed models describing the nonmotile consortia engaged in metabolic exchange^10,13^. Detailed procedures for model construction, simulation setup, and analytical protocols are provided in the Supplementary Information. Definitions and default values of all variables and parameters are summarized in Table S2. The individual-based simulations were implemented in Wolfram Mathematica (Version 14.0), and the scripts are available online (GitHub: https://github.com/WMXgg/Cell-motility-project-codes/tree/main/PA%20project%20Code/Simcodes).

### Quantification and Statistical Analysis

All statistical analyses were performed in Wolfram Mathematica (Version 14.0) using built-in statistical functions. Student’s t-tests were conducted with the TTest function, and significance levels are indicated in the corresponding figure panels. Linear regressions were performed using the LinearModelFit function (Wolfram Mathematica, Version 12.0), and the corresponding adjusted R^2^ values are reported in all relevant figures. Spearman’s rank correlations were computed with the SpearmanRankCorrelation function. The quantitative indices, including the intermixing index, Moran’s I, *Tm, gE, T*_*cess*_, and *T*_*agg*_, were calculated using custom scripts written in Wolfram Mathematica. Details regarding replicate numbers and the statistical tests applied are provided in the respective figure legends.

To compare the fitted correlations between single-cell growth rate and local partner fraction across the motile and nonmotile consortia, we applied a three-step statistical framework. First, we performed a Chow Test^90^ (equivalent to a likelihood ratio test for nested linear models) to assess the null hypothesis that the two datasets follow the same linear regression model. This test compares the residual sum of squares (SSEp) of a pooled regression to those (SSEm and SSEnm) of separate regressions fitted to motile and nonmotile cells, yielding an *F*-statistic calculated as

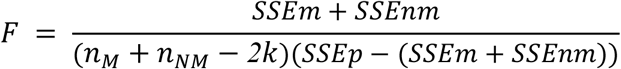

where *k* = 2 is the number of model parameters, *n*_*M*_ and *n*_*NM*_ denote the sample sizes of the motile cells and nonmotile cells. *F* follows an *F*-distribution with degrees of freedom (k, *n*_*M*_ + *n*_*NM*_ *– 2k*). The corresponding *p*-value was calculated to determine whether the two datasets showed a statistically significant departure from a common regression model. Second, a regression-with-indicators approach was used to determine whether the difference was driven primarily by intercepts, slopes, or both. Data from both treatments were pooled and fitted using a linear model with a binary group indicator and an interaction term between group and partner fraction:

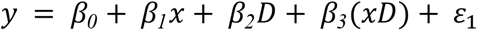

This parameterization enables direct hypothesis tests for parameter homogeneity across groups: a significant *β*_*3*_ indicates that the slope differs between groups, while a significant *β*_*2*_ indicates an intercept shift between groups at a common slope. Statistical significance was assessed using Student’s t-tests. Third, to quantify the magnitude of similarity between the fitted relationships, a Model Difference Index (MDI) was defined as the maximum of the mean point-to-line distance (*D*_*M*→*NM*_ or *D*_*NM*→*M*_) across the two datasets, divided by the corresponding within-dataset RMS residual (σ_*1*_ or σ_*2*_):

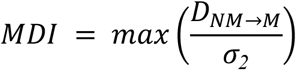

Intuitively, MDI measures the systematic deviation between two fitted lines compared with the typical scatter of the data around each line. The observed MDI values were compared to a Null Model. Specifically, for each dataset, the original x-values were held fixed, and synthetic y-values were generated according to a linear model of the form

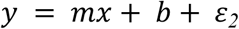

where the slope *m* and intercept *b* were randomly sampled from distributions centered on the values estimated from the original data. The noise term *ε*_*2*_ was drawn from a zero-mean normal distribution, with its variance chosen to yield a target coefficient of determination within a predefined range (0.3 to 0.9), ensuring that the generated datasets exhibited comparable linear strength to the observed data. For each null realization, linear regression was performed independently for the two synthetic datasets, and the corresponding MDI was computed. A total of 5,000 null realizations were generated per comparison, yielding an empirical null distribution against which the observed MDI was evaluated.

## Author Contributions

W.M.X. and M.A. conceived the study. W.M.X. constructed the bacterial strains, performed the microfluidic experiments, conducted image analyses, and designed and implemented the individual-based modeling. W.M.X. wrote the manuscript with input from M.A. and O.S.. W.M.X., O.S., and M.A. acquired the funding. O.S. and M.A. supervised the research.

## Acknowledgements

We thank Dr. Luyan Z. Ma (Institute of Microbiology, Chinese Academy of Sciences) for kindly providing the *Pseudomonas aeruginosa* PAO1 mutant strains. We are grateful to the late Dr. Alma Dal Co for her valuable suggestions on the conceptualization and development of the microfluidic methods used in this study. We also thank Dr. Simon van Vliet for his helpful advice on culturing *P. aeruginosa* in monolayer microfluidic devices. We further acknowledge the members of the Microbial Systems Ecology group for insightful discussions throughout this work. Miaoxiao Wang sincerely thanks his wife, Yaxi Li, and their precious baby, Ximing Wang, for their unwavering love and support throughout this project. This research was supported by a National Key R&D Program of China (grant number 2025YFA0921700 to W.M.X.), a Joint Research Project grant from the Swiss National Science Foundation (grant no. IZLCZ0_206044 to W.M.X. and M.A.), the National Natural Science Foundation of China (grant no. 32500081 to W.M.X.), and as part of the NCCR Microbiomes, funded by the Swiss National Science Foundation (grant nos. 51NF40_180575 and 51NF40_225148 to O.S. and M.A.)., and by ETH Zurich and Eawag.

