## Supplementary Information for "Cell motility enhances metabolic coupling in spatially structured microbial communities"

**S1 Model Description**

Here, we describe the model framework in detail. The model is a spatially explicit, individual-based simulation implemented on a 40 × 40 lattice, adapted from our previous work^1,2^. The simulation package was developed in Wolfram Mathematica (Version 14.0) and executed on a Windows-based cloud server (https://www.yisu.com/). All model parameters and their values are summarized in Table S2. The scripts are available online: <https://github.com/WMXgg/Cell-motility-project-codes/tree/main/PA%20project%20Code/Simcodes>.

**S1.1 Equations characterizing metabolic exchanges and microbial growth**

The individual-based model considers two interacting cell types, denoted as type A and type B. Type A produces only amino acid 1, whereas type B produces only amino acid. Consequently, the growth of type A depends on the supply of amino acid 2 leaked by type B, and vice versa. Cells were modeled as spherical objects positioned on a 40 × 40 lattice with rigid boundaries, corresponding to a two-dimensional regular grid. The morphology of space and the maximum number of cells in the lattice (1600) matched that observed in our 40-μm square microfluidic chambers. The internal (I) and external (E) concentrations of amino acids were also modeled dynamically in two dimensions. Metabolic exchanges and microbial growth were modelled following six key assumptions and using functions identical to our previous study^1^:

1) Cells actively take up amino acids from the external environment following a first-order kinetic with uptake rate *r^u^_i_*, *uptake* = *r^u^_i_* · *E_i_*.
2) Cells leak amino acids passively with a leakage rate *r^l^_i_*, *leakage* = *r^l^_i_* · (*I_i_* − *E_i_*).
3) Cells consume amino acids proportionally to their growth rate, *consumption* = *µ* · *I_i_* .
4) Cell growth is limited by the amino acid that they cannot produce, following a Monod kinetic: *µ* = (*µ^aux^_i_* ·*I_i_*) / (*K_i_*+*I_i_*).
5) Cells produce amino acids. Cells maintain a constant concentration *I^c^_i_* of the amino acid that they can produce as a result of tight regulation of the amino acid production rate^3,4^.
6) Amino acids diffuse in the extracellular environment with an effective diffusion coefficient that depends on local cell density: *D_effi_* = (1−*c*)·*D_i_* / (1+ *c*/2 ), where *D_i_* is the diffusion constant in the empty space and *c* is the cell density.

In each lattice grid occupied by a cell, the temporal dynamics of the internal (*I*) and external (*E*) concentrations of amino acids are modeled as follows (the subscript *i* denotes amino acid 1 or 2):

$\frac{\text{∂}\text{I}_{\text{1}}}{\text{∂t}}\text{ }\text{=}\text{ }\text{0}$	if type is A	[1]

$\frac{\text{∂}\text{I}_{\text{1}}}{\text{∂t}}\text{ = }\text{r}\text{u}\text{1}\text{ · }\text{E}\text{1}-\text{r}\text{l}\text{1}\text{ }\text{· (}\text{I}\text{1}\text{ - }\text{E}\text{1}\text{)}-\frac{{\text{µ}^{\text{aux}}}_{\text{1}}\text{ ·I}\text{1}}{\text{(K}\text{1}\text{+I}\text{1}\text{)}}\text{·I}\text{1}$	if type is B	[2]

$\frac{\text{∂}\text{I}_{\text{2}}}{\text{∂t}}\text{ = }\text{r}\text{u}\text{2}\text{ · }\text{E}\text{2}-\text{r}\text{l}\text{2}\text{ }\text{· (}\text{I}\text{2}\text{ - }\text{E}\text{2}\text{)}-\frac{{\text{µ}^{\text{aux}}}_{\text{2}}\text{ ·I}\text{2}}{\text{(K}\text{2}\text{+I}\text{2}\text{)}}\text{·I}\text{2}$	if type is A	[3]

$\frac{\text{∂}\text{I}_{\text{2}}}{\text{∂t}}\text{ }\text{=}\text{ }\text{0}$	if type is B	[4]

$\frac{\text{∂}\text{E}_{\text{i}}}{\text{∂t}}\text{ =}\text{-α·}\text{ }\text{r}\text{u}\text{i}\text{ · }\text{E}_{\text{i}}-\alpha\cdot\text{r}\text{l}\text{i }\text{· (}\text{I}\text{i}\text{ - }\text{E}\text{i}\text{)}-\text{D}_{\text{effi}}\text{∙}\text{∇}^{\text{2}}\text{E}\text{i}$	[5]

Where α = *V_in_* / *V_out_* = *c* / (1 − *c*) represents the ratio between the intracellular (*V_in_*) and extracellular (*V_out_*) volume. To reduce the number of model parameters, we nondimensionalized the amino acid concentrations by expressing them relative to their Monod constants:

$\text{φ}_{\text{i}}\text{=}\frac{\text{I}_{\text{i}}}{\text{K}\text{i}}$,	$\text{ϵ}_{\text{i}}\text{=}\frac{\text{E}_{\text{i}}}{\text{K}\text{i}}$

Substituting these variables yields the following nondimensional equations:

$\frac{\text{∂}\text{φ}_{\text{1}}}{\text{∂t}}\text{ }\text{=}\text{ }\text{0}$	if type is A	[6]

$\frac{\text{∂}\text{φ}_{\text{1}}}{\text{∂t}}\text{ = }\text{r}\text{u}\text{1}\text{ · }\text{ϵ}\text{1}-\text{r}\text{l}\text{1}\text{ }\text{· (}\text{φ}\text{1}\text{ - }\text{ϵ}\text{1}\text{)}-\frac{{\text{µ}^{\text{aux}}}_{\text{1}}\text{ ·φ}\text{1}}{\text{(}\text{1}\text{+φ}\text{1}\text{)}}\text{·φ}\text{1}$	if type is B	[7]

$\frac{\text{∂}\text{φ}_{\text{2}}}{\text{∂t}}\text{ = }\text{r}\text{u}\text{2}\text{ · }\text{ϵ}\text{2}-\text{r}\text{l}\text{2}\text{ }\text{· (}\text{φ}\text{2}\text{ - }\text{ϵ}\text{2}\text{)}-\frac{{\text{µ}^{\text{aux}}}_{\text{2}}\text{ ·φ}\text{2}}{\text{(}\text{1}\text{+φ}\text{2}\text{)}}\text{·φ}\text{2}$	if type is A	[8]

$\frac{\text{∂}\text{φ}_{\text{2}}}{\text{∂t}}\text{ }\text{=}\text{ }\text{0}$	if type is B	[9]

$\frac{\text{∂}\text{ϵ}_{\text{i}}}{\text{∂t}}\text{ =}\text{-α·}\text{ }\text{r}\text{u}\text{i}\text{ · }\text{ϵ}_{\text{i}}-\alpha\cdot\text{r}\text{l}\text{i }\text{· (}\text{φ}\text{i}\text{ - }\text{ϵ}\text{i}\text{)}-\text{D}_{\text{effi}}\text{∙}\text{∇}^{\text{2}}\text{ϵ}\text{i}$	[10]

Since the primary goal of this model was to investigate the effects of cell motility, we further simplified it by assuming that the two cell types share identical growth and interaction parameters:

$\text{r}\text{u}\text{1}\text{ }\text{=}\text{ }\text{r}\text{u}\text{2}\text{ }\text{=}\text{ rU}$, $\text{r}\text{l}\text{1}\text{ }\text{=}\text{ }\text{r}\text{l}\text{2}\text{ }\text{=}\text{ rL}$, ${\text{µ}^{\text{aux}}}_{\text{1}}\text{ }\text{=}\text{ }{\text{µ}^{\text{aux}}}_{\text{2}}\text{ }\text{=}\text{ }\text{g}\text{E}$, $\text{D}_{\text{1}}\text{ }\text{=}\text{ }\text{D}_{\text{2}}\text{ }\text{=}\text{ }\text{D}$

Therefore, only these four parameters were directly related to the growth and metabolic interactions, and their effects on the benefits of the motility are thoroughly examined.

**S1.2 Diffusion of the amino acids**

The diffusion of amino acids was modeled using a second-order approximation for two-dimensional diffusion, consistent with our previous work^2^. In this approach, the diffusion rate between neighboring lattice elements is proportional to the concentration gradient of the amino acid. For each lattice grid and diffusion step, we computed the weighted mean concentration of the amino acid within its 9-element Moore neighborhood, denoted as $\underline{\epsilon}_{i}$. The diffusive flux across grid boxes was then estimated using the term $D_{effi}\cdot\underline{\epsilon}_{i}$ , which represents the diffusion component $D_{effi}\cdot\nabla^{2}\epsilon i$ in Equation [10].

**S1.3 Cell division**

Each cell was initialized with a biomass *X_0_*. When a cell’s biomass reached an upper threshold, *X*_max_ = (2+*ε*) *X_0_*, it divided into two daughter cells of equal biomass. Here, *ε* represents stochastic noise in the cell cycle, modeled as a normally distributed random variable with a mean of 0 and standard deviation (0, *θ*). Following division, one cell remained in its original grid position, while the other cell was randomly assigned to one of the eight directly adjacent grids. If the target grid was already occupied, the daughter cell competed with the resident cell for occupancy; one of the two cells was then randomly removed with equal probability (0.5).

**S1.4 Assumptions of cell motility**

We incorporated three distinct motility patterns into our model:

(1) Nonmotile state, where cells remain stationary;

(2) Random motility, representing purely stochastic movement that serves as a null model;

(3) Aggregative motility, representing movement guided by a QS signal.

Below, we describe in detail how random and aggregative motility were implemented in the model.

*S1.1 Assumptions of random motility*

Random motility was modeled using the classical *run–tumble* framework^5,6^. In each time step (iteration), the behavior of a motile cell was simulated through the following three stages:

(1) **Movement decision**: The model first determines whether the cell will move, governed by a *movement probability* (*Mp*).

(2) **Cell reorientation**: If the cell moves, its orientation may change based on a *reorientation noise* parameter (*Rn*). The turning angle is drawn from a normal distribution (0, *Rn*). A higher *Rn* value corresponds to greater randomness and larger turning angles, leading to more diffusive trajectories.

(3) **Displacement**: The distance that the cell move within one time step follows a normal distribution (*Ms*×*dt*, 1), where *Ms* represents the speed of the movement and *dt* represents the time step.

To account for physical constraints, cells were assumed to reverse their movement direction upon colliding with rigid chamber boundaries or encountering another occupied lattice site^7^. This rule prevented cell overlapping and mimicked volume exclusion effects observed in densely packed microbial populations.

*S1.2 Assumptions of aggregative motility*

Aggregative motility was modeled as an extension of the random motility framework by incorporating quorum-sensing (QS)-mediated behavioral control. We assumed that all cells continuously secrete a diffusible QS signal that accumulates in the extracellular environment. At each simulation step, every cell senses the local concentration of this signal within its grid. When the sensed concentration exceeds a predefined threshold (*St*), the cell ceases movement for that iteration. This implementation captures the self-reinforcing “rich-get-richer” dynamics characteristic of signal-guided twitching motility observed in Pseudomonas aeruginosa^8,9^, where cells preferentially accumulate and remain within high-signal regions. Consequently, this mechanism leads to the spontaneous formation of central aggregates, mimicking biologically observed clustering patterns driven by QS-controlled motility regulation.

**S1.5 Simulation processing**

The model was implemented on a 40 × 40 lattice, with each grid cell accommodating only a single biological cell. At the start of each simulation, the extracellular amino acid concentrations were initialized to zero across all grids. Cells were randomly assigned to grid positions according to specified initial frequencies, each with an initial biomass *X_0_*. Simulations terminated once 95% of the grids were occupied or when 10,000 iteration steps had elapsed. The simulation framework was integrated into a custom-built package developed in Wolfram Mathematica (Version 14.0), which included several dedicated functions:

(1) leakFun — computes amino acid leakage;

(2) biopubChange — updates amino acid uptake and cell growth;

(3) cellReproduce — simulates cell division events;

(4) nutpubSpread — simulates amino acid diffusion;

(5) Move — simulates random motility;

(6) MoveQS — simulates quorum-sensing–mediated aggregative motility.

For the nonmotile consortium, functions (1)–(4) were executed. For the randomly motile consortium, functions (1)–(5) were applied, whereas the aggregatively motile consortium utilized functions (1)–(4) and (6). Although these biological processes occur simultaneously in nature, computational constraints require sequential execution of these processes as well as the lattice grids. To minimize order-related bias, the simulation order of these processes and the sequence of grid computation were randomized at each iteration.

For simulations using specific parameter sets, parameters were manually defined within the Parameter.m script. For large-scale simulations involving diverse parameter sets, the parameter sets were randomly generated using the RandomReal function of Wolfram Mathematica, with the range of each parameter defined in Table S2. All these parameter sets were stored in several .csv files and automatically imported by the Mathematica package, enabling high-throughput processing. Simulations were performed on a Windows-based cloud server (https://www.yisu.com/). The complete simulation package is publicly available on GitHub: <https://github.com/WMXgg/Cell-motility-project-codes/tree/main/PA%20project%20Code/Simcodes/Simulation%20Package>.

**S1.6 Quantification of model outputs**

During each simulation, key variables—including cell positions, biomass, growth rates, and intra- and extracellular amino acid concentrations—were recorded every 10 iteration steps. These data were subsequently used to calculate defined quantitative indices and to perform fitting analyses.

All quantitative indices—including the intermixing index, Moran’s I, *Tm*, *gE*, *T_cess_*, and *T_agg_*—were computed using custom scripts written in Wolfram Mathematica (Version 14.0). The intermixing index was defined in the same way as in the experimental analyses. Specifically, for each type-A cell, we counted the number of type-B and type-A cells located within a fixed neighborhood of five grid units (equivalent to approximately 5 μm in the experimental definition) and calculated the ratio between the number of type-B and type-A cells. The overall intermixing index at a given time point was obtained as the mean of these ratios across all type-A cells. Definitions and computational details of other indices are provided in the corresponding figure legends.

All statistical analyses of the simulation outputs were performed in Wolfram Mathematica (Version 14.0) using built-in statistical functions. Student’s t-tests were performed using the TTest function, and significance levels are indicated in the figure panels. Linear regressions were fitted using the LinearModelFit function (Version 12.0), with adjusted R^2^ values reported in the corresponding figures. Spearman’s rank correlations were calculated using the SpearmanRankCorrelation function. The number of replicates and the specific statistical tests used are detailed in each figure legend.

### S2 Supplementary Tables

**Table S1 |** Statistical comparisons of the linear regression models fitted in Figure 3e-f and Figure S6

| Chamber geometries | genotypes | Chow Test *p*-values | Slope *p*-values | Intercept *p*-values | MDI | Similarity percentile in the Null model |
| --- | --- | --- | --- | --- | --- | --- |
| 60 μm circular | Δ*hisD* | < 10e-6 | < 10e-6 | < 10e-6 | 1.19 | 0.911 |
|  | Δ*trpB* | < 10e-6 | 0.490 | 6.44e-4 | 0.986 | 0.980 |
| 40 μm circular | Δ*hisD* | < 10e-6 | 1.62e-4 | < 10e-6 | 0.995 | 0.981 |
|  | Δ*trpB* | < 10e-6 | 0.42 | < 10e-6 | 0.985 | 0.982 |
| 60 μm square | Δ*hisD* | < 10e-6 | 1.62e-5 | < 10e-6 | 1.01 | 0.975 |
|  | Δ*trpB* | 1.21e-6 | 0.0824 | 1.41e-6 | 1.00 | 0.972 |
| 40 μm square | Δ*hisD* | < 10e-6 | < 10e-6 | < 10e-6 | 1.08 | 0.951 |
|  | Δ*trpB* | < 10e-6 | 0.441 | 1.06e-6 | 0.98 | 0.983 |

**Table S2 |** Model parameters, default values, and the test values ranges used in the individual-based simulations

| Symbol | Parameter description | Unit | Default value | Range tested | References |
| --- | --- | --- | --- | --- | --- |
| $\partial t$ | iteration step | min | 1 | – | N.A. |
| *lenGrid* | Side length of the simulation space | grid | 40 | 20, 40, 60, 80, 100 | N.A. |
| *ID* | Initial cell density | dimensionless | 0.025 | 0.005, 0.01, 0.025, 0.05, 0.1 | N.A. |
| *IB* | Initial cell biomass | fg | 150 | – | ^2^ |
| *Bl* | Biomass limit for cell division | fg | 300 | – | ^2^ |
| *Bn* | Noise term for biomass (bionoise) | fg | 30 | – | ^2^ |
| *Mp* | Maximum movement probability | dimensionless | 0.9 | 0.1 – 0.9 | N.A. |
| *Ms* | Mean value of moving speed | grid (μm)/min | 1 | 10^-0.5^ – 10^0.5^ | ^6,7^ |
| *Rn* | Rotation noise (SD of reorientation angle); higher Rp indicates greater randomness of the rotation. | rad | 1 | 10^-0.5^ – 10^0.5^ | ^6,7^ |
| *St* | Threshold of the signal concentration for stopping movement | – | 100 | 10^0^ – 10^3^ | N.A. |
| *D* | Diffusion coefficient of amino acids | μm²/min | 1 | 10⁻¹ – 10⁰ | ^10–12^ |
| *rL* | Leakage rate of amino acids | /min | 10⁻³ | 10⁻⁵ – 10⁻¹ | ^1,13^ |
| *rU* | Uptake rate of amino acids | /min | 10² | 10⁻¹ – 10⁴ | ^14–16^ |
| *gE* | Essential (maximum) growth rate | /min | 0.1 | 10⁻³ – 10⁰ | ^17,18^ |

### S3 Supplementary Figures

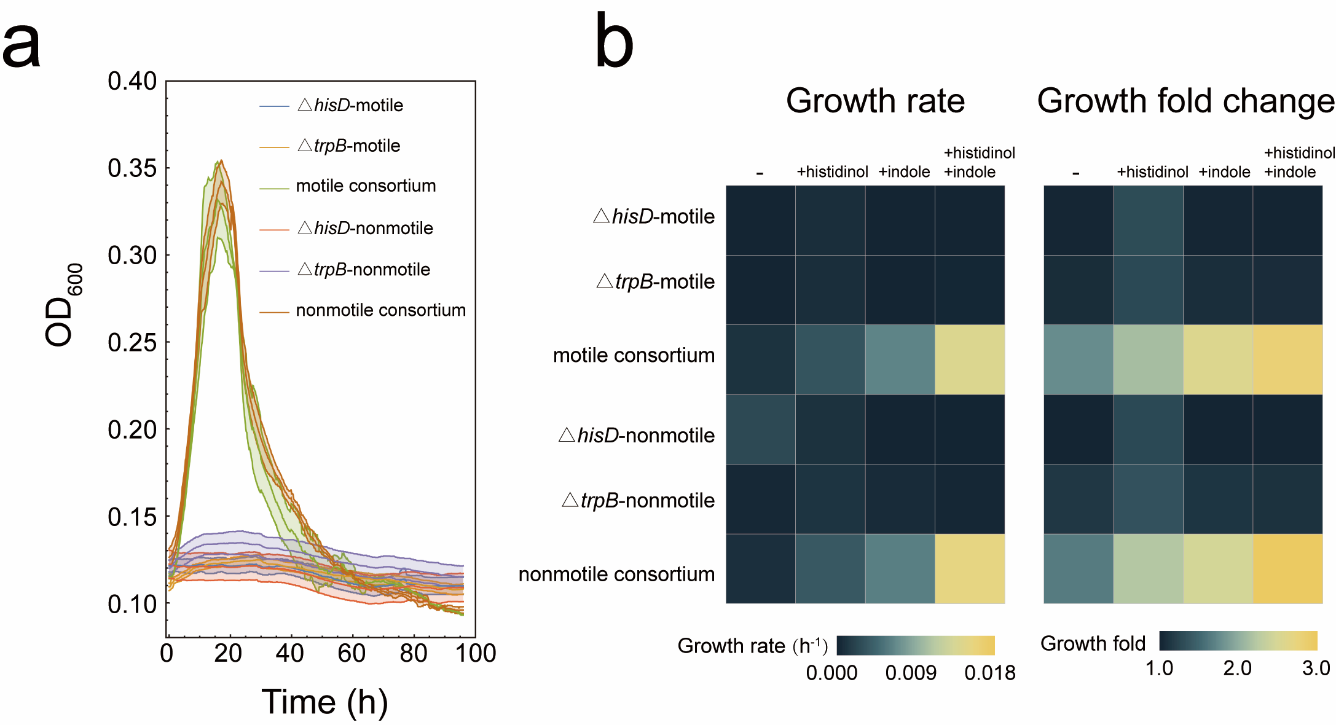

**Figure S1 | Liquid monocultures and cocultures of the auxotrophs used in this study.**

Growth analyses of the motile and non-motile auxotrophs cultured in 96-well plates in defined minimal medium with or without supplementation of the amino acid precursors histidinol and/or indole. Optical density at 600 nm (OD_600_) was tracked over time. (a) Growth curves. (b) Growth rates (left) and fold changes of (right) were obtained by fitting the Gompertz function to derive growth rates and fold-changes. All conditions were tested with six independent replicates.

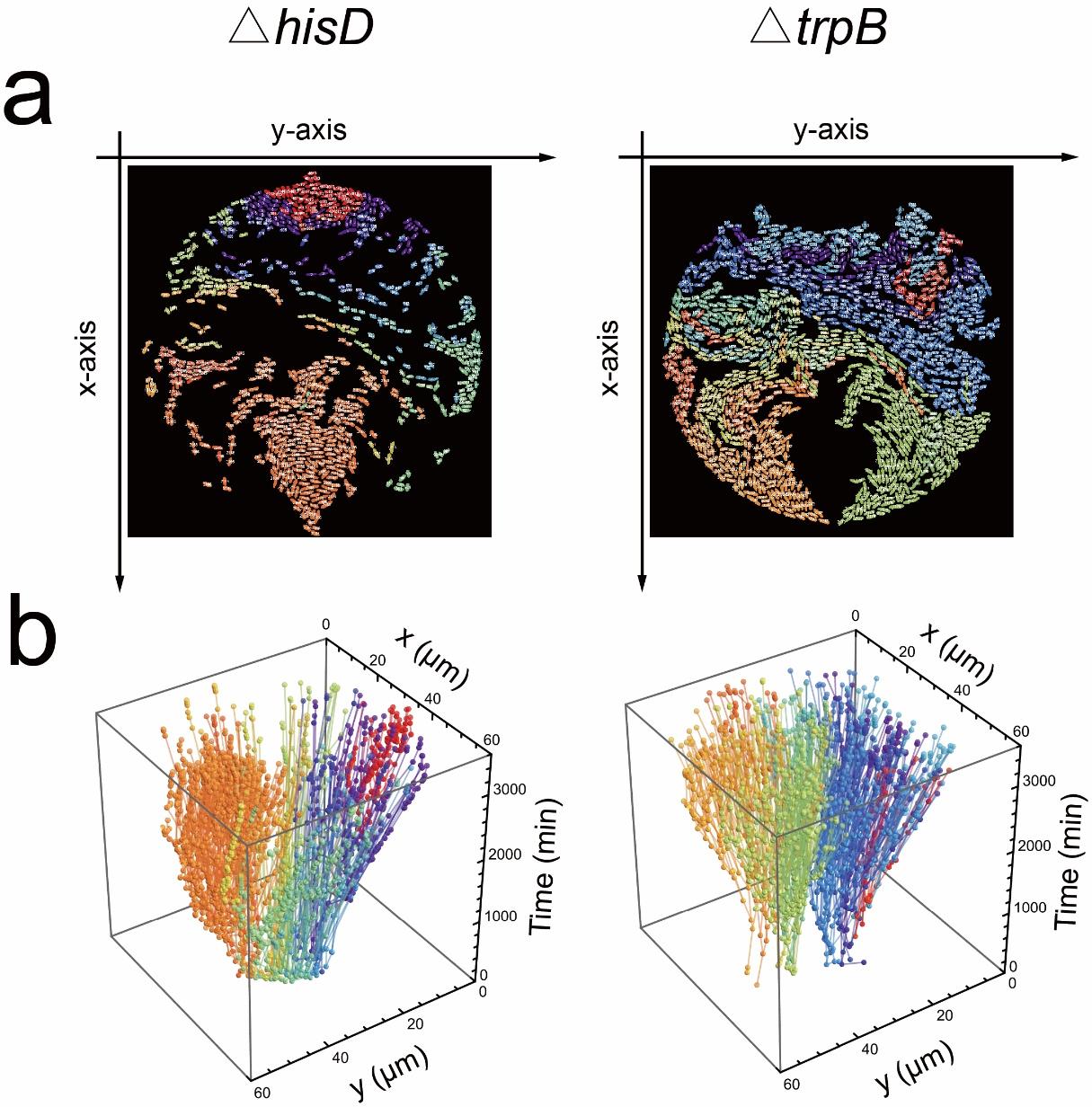

**Figure S2 | Lineage tracking of the nonmotile consortium.**

(a) Microscopic images of the nonmotile consortium grown in microfluidic chambers, showing histidine auxotrophs (left) and tryptophan auxotrophs (right). Cells derived from the same founder are assigned identical colors and number codes based on lineage tracking across divisions. The final state at 59.5 h is shown. Original time-lapse images are provided in Supplementary Video 1 (right), and corresponding time-lapse lineage tracking in Supplementary Video 5. (b) Reconstructed lineage trees from time-lapse images. Cells (spheres, colored as in (a)) are plotted according to their spatial coordinates (x, y) and time (t). Lines connect parent and progeny cells, with branching points indicating division events.

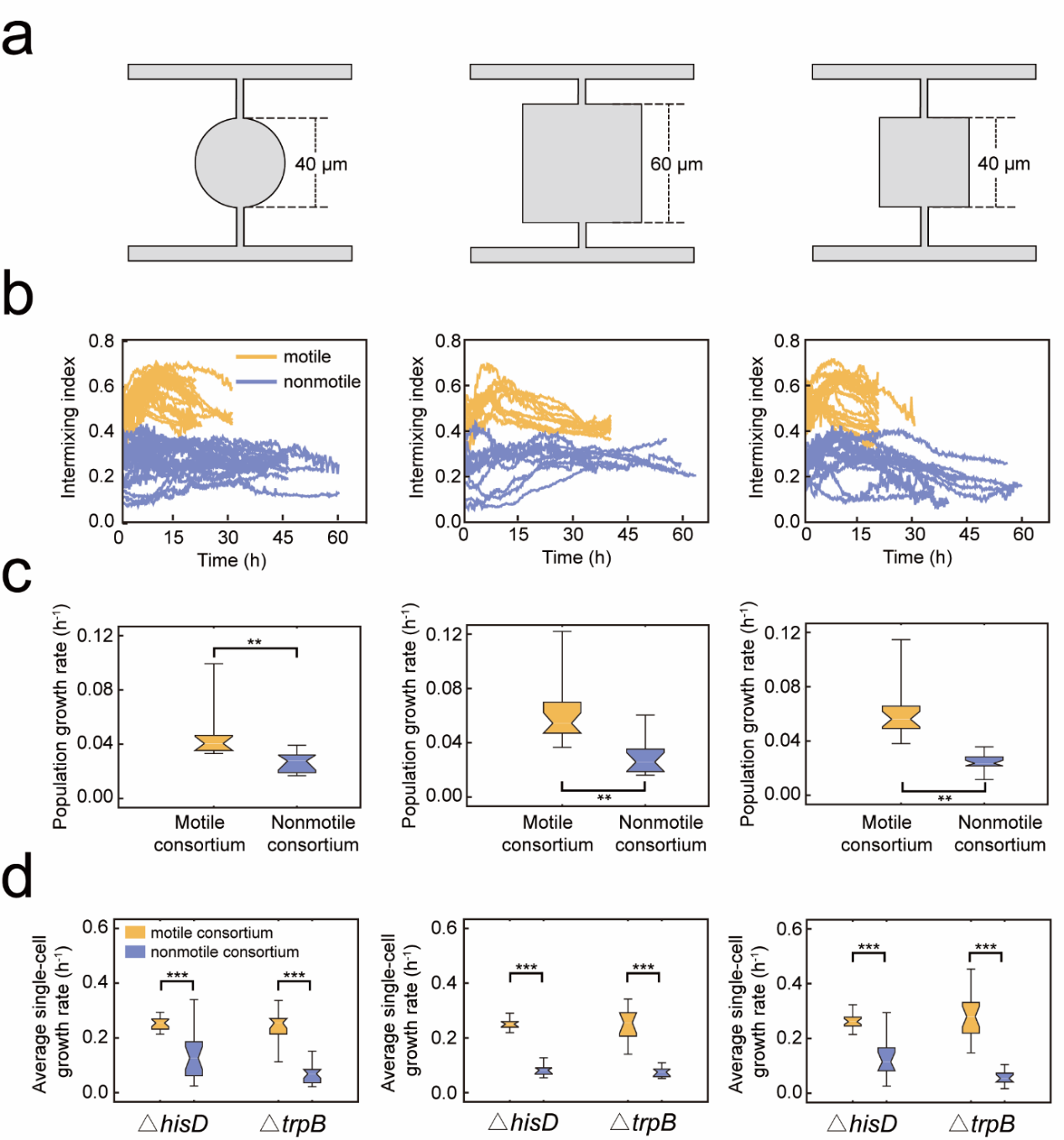

**Figure S3 | Quantitative analyses in microfluidic chambers of three different geometries.**

(a) Schematics of the three-chamber designs. (b–e) Comparison between motile and non-motile consortia in terms of intermixing index (b), population-level growth rates (c), and average single-cell growth rates (d). Data represent replicates from three independent experiments: 21 motile and 19 nonmotile replicates in 40-μm diameter circular chambers; 12 motile and 8 nonmotile replicates in 60-μm square chambers; and 22 motile and 14 nonmotile replicates in 40-μm square chambers. Statistical significance was assessed by Student’s t-test: *: *p* < 0.05; **: *p* < 0.01; ***: *p* < 0.001.

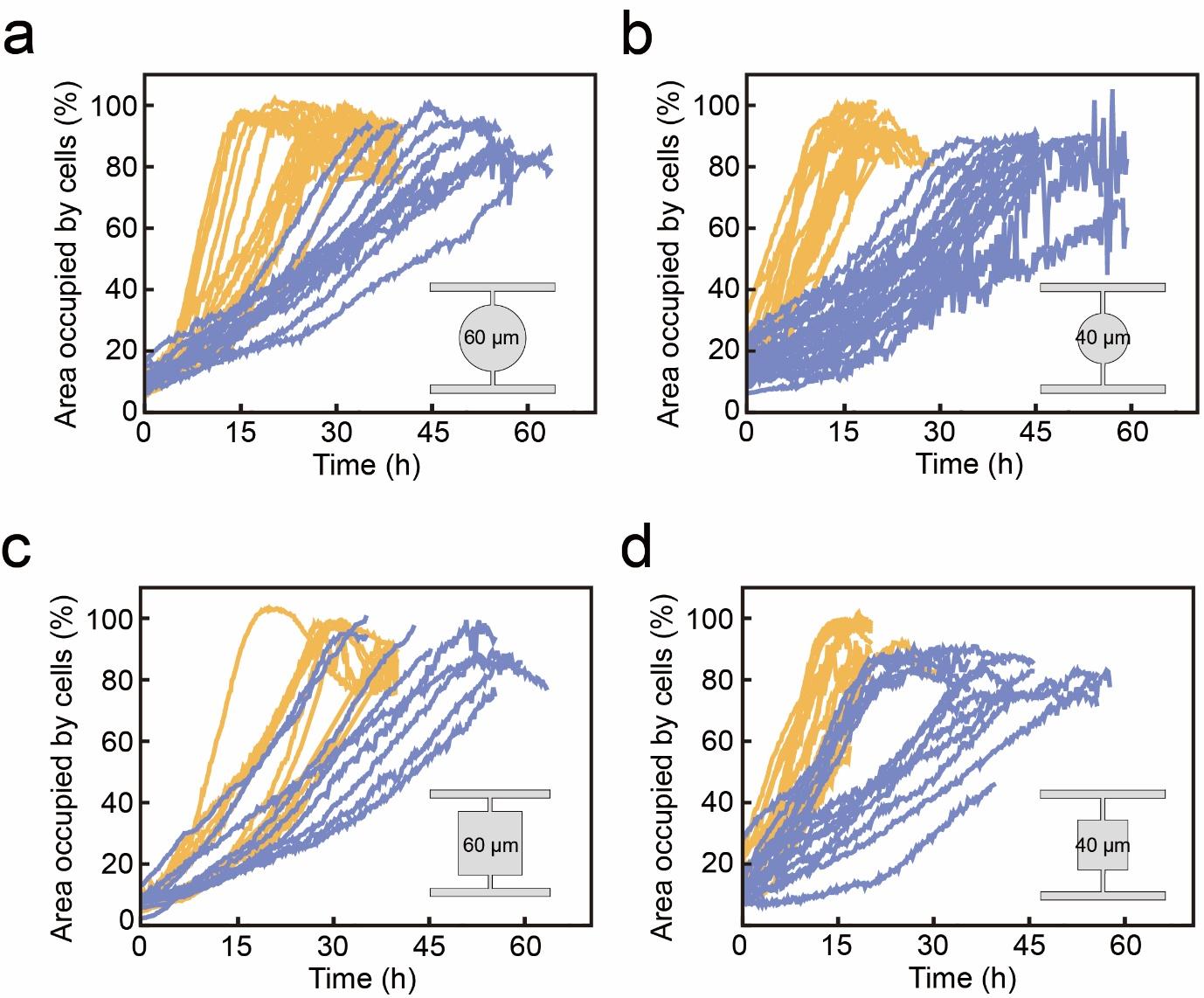

**Figure S4 | Growth dynamics of motile and nonmotile consortia across chamber geometries.**

Growth curves of motile and nonmotile consortia in 60-μm diameter circular chambers (a), 40-μm diameter circular chambers (b), 60-μm square chambers (c), and 40-μm square chambers (d). Growth was quantified as the fraction of chamber area occupied by cells, measured from segmented time-lapse images.

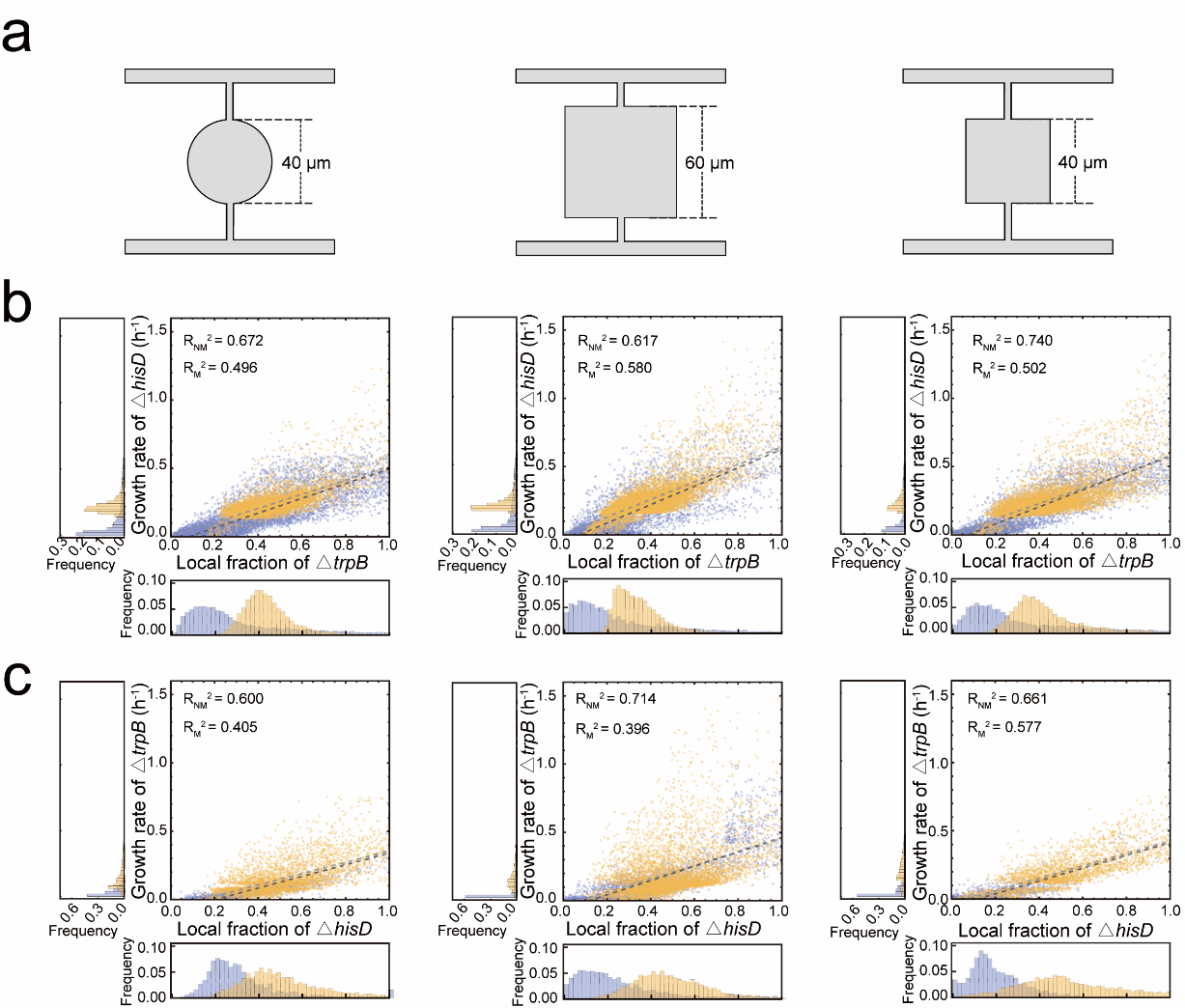

**Figure S5 | Correlation between the single-cell growth rates and the fraction of the partner within the local range.**

(a) Schematics of the three-chamber designs. (b-c) Histidine (b) and tryptophan (c) auxotrophic cells in both the motile and nonmotile consortium grew faster with an increasing fraction of the partner within the local range (5 μm). Yellow dots represent single cells in the motile consortium (12,891, 29,847, and 19,779 Δ*hisD* cells, as well as 7,429, 10,309, and 8,113 Δ*trpB* cells for the three chamber designs, respectively), while blue dots represent single cells in the nonmotile consortium consortium (19,662, 14011, and 12,277 Δ*hisD* cells, as well as 7,859, 11,188, and 4,864 Δ*trpB* cells for the three chamber designs, respectively). The dark grey line represents the linear regression of the nonmotile cells with the coefficient of determination denoted as R^2^_NM_, while the dark grey line represents the linear regression of the motile cells with the coefficient of determination denoted as R^2^_M_. The histograms denote the frequency distributions of the local partner frequency and the single-cell growth rates.

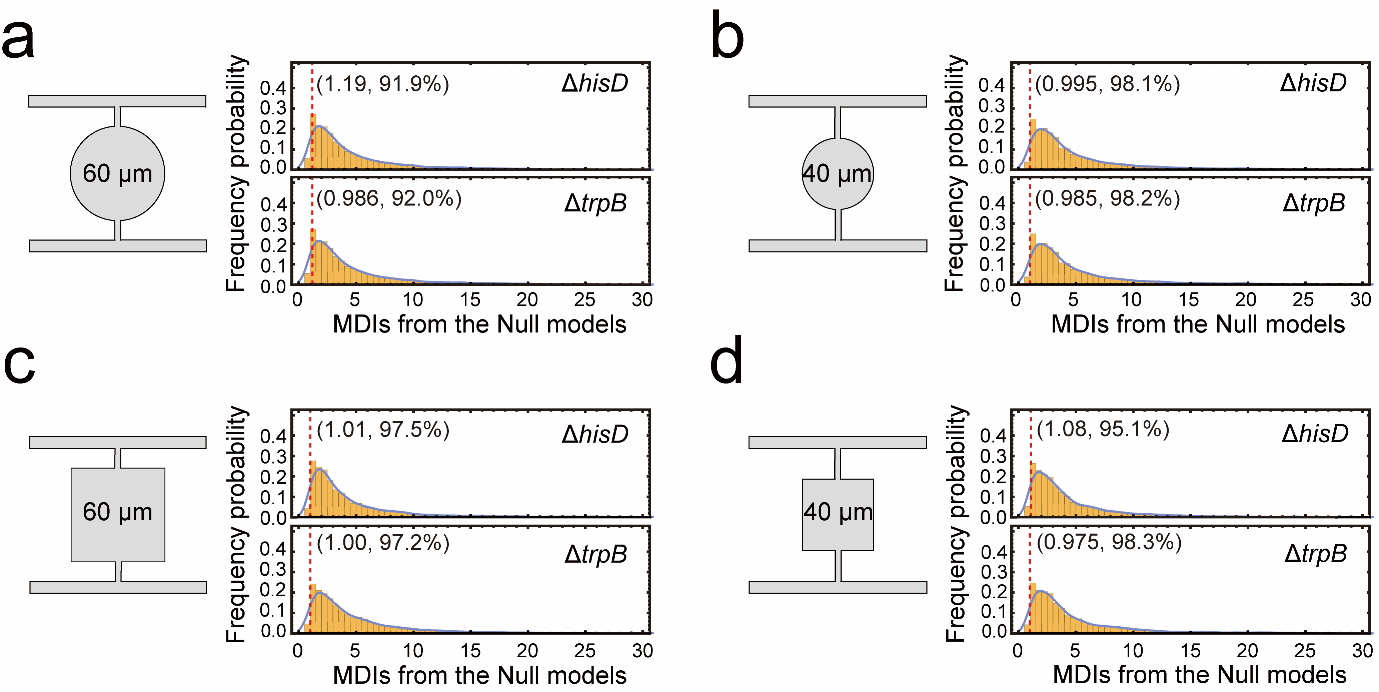
**Figure S6 | Null model analyses quantifying the similarity in the fitted relationships linking single-cell growth rate and local partner fraction between the motile and nonmotile consortia.**

Histograms show the null distributions of the Model Difference Index (MDI), with blue curves indicating kernel density estimates. MDI is defined as the maximum mean point-to-line distance between datasets, normalized by the within-dataset RMS residual. The vertical red line marks the observed MDI value (MDI_observed_). The MDI_observed_ and the associated percentile values, Pr(MDI_Null_ ≥ MDI_observed_), are indicated next to the red line. Analyses are shown for four chamber geometries (panels a–d) and for both auxotrophic genotypes. 5000 null model samples were generated for each comparison. Details of the MDI calculation and null-model construction are provided in the Methods section.

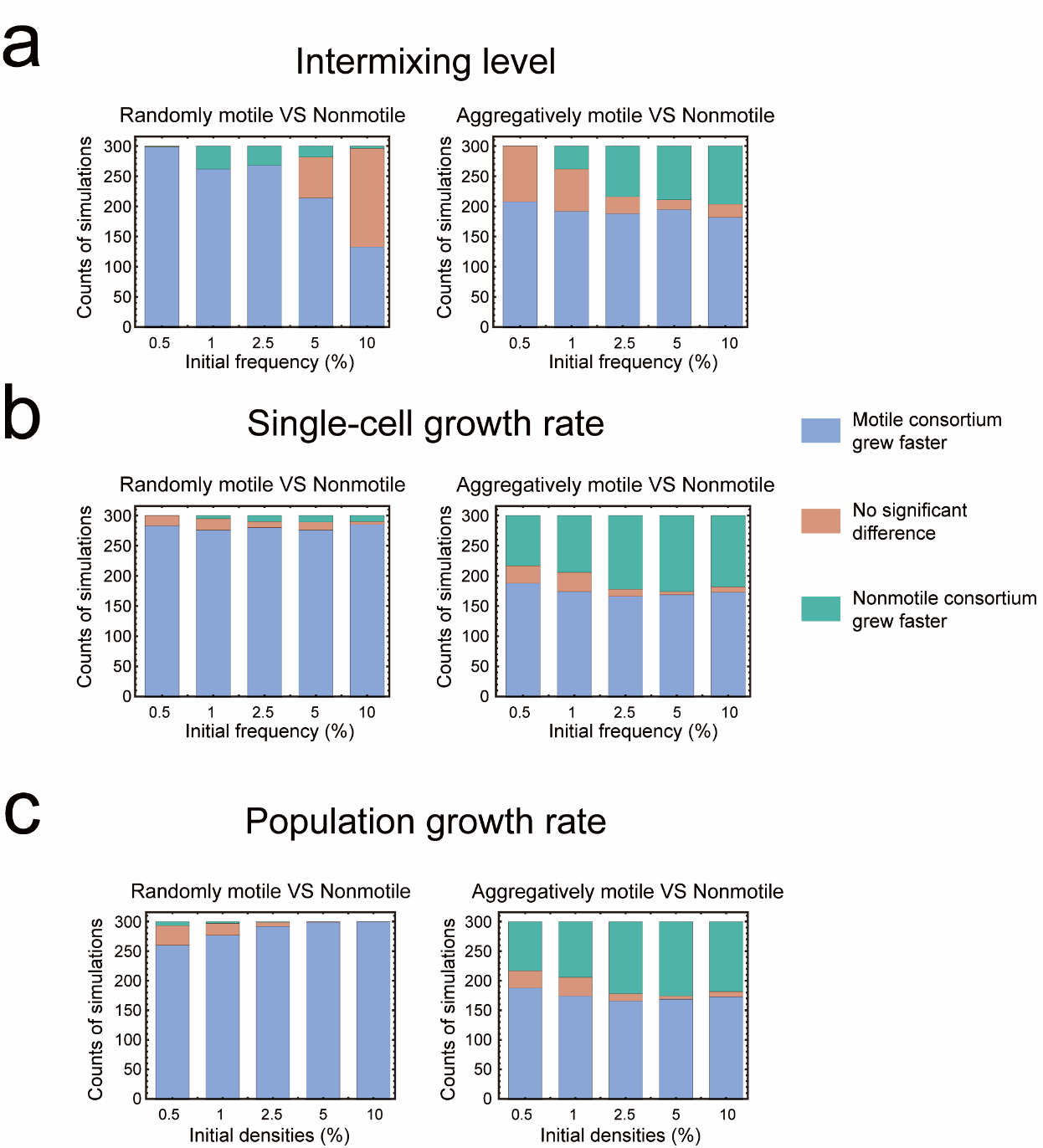

**Figure S7 | Effects of initial cell density on the positive effects of motility.**

Summary of whether motile consortia exhibited higher intermixing levels (a), single-cell growth rates (b), and population growth rates (c) compared with nonmotile consortia under five different initial cell densities. Statistical significance was assessed at *p* < 0.05.

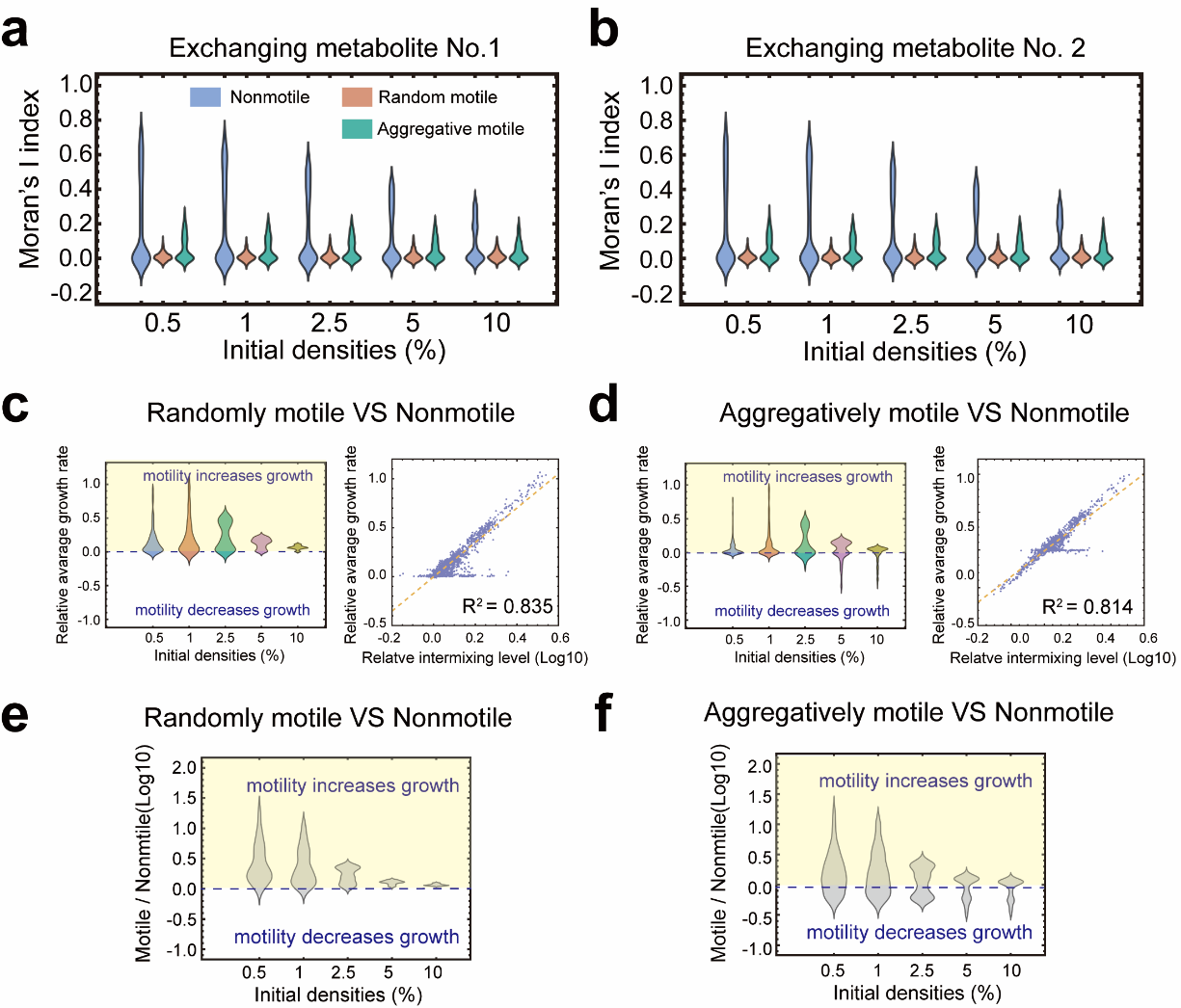

**Figure S8 | Comparison of metabolite distribution and single-cell growth rates between the motile and nonmotile consortia.**

(a–b) Evenness of metabolite distributions across five initial cell densities, quantified using Moran’s I index. Moran’s I values range from -1 to 1. Values close to 0 indicate a spatially even distribution of metabolites. Positive values approaching 1 indicate strong spatial clustering, where neighbouring grids hold similar metabolite concentrations. Negative values approaching -1 indicate a dispersed or checkerboard-like pattern, where neighbouring grids differ strongly in the concentrations of metabolites. (c–d) Relative average single-cell growth rates of randomly motile (c) and aggregately motile (d) consortia versus nonmotile consortia. Left: value distributions across five initial densities; right: correlations between relative growth rates and intermixing level. (e–f) Relative population growth rates of randomly motile (e) and aggregately motile (f) consortia versus nonmotile consortia across five initial cell densities.

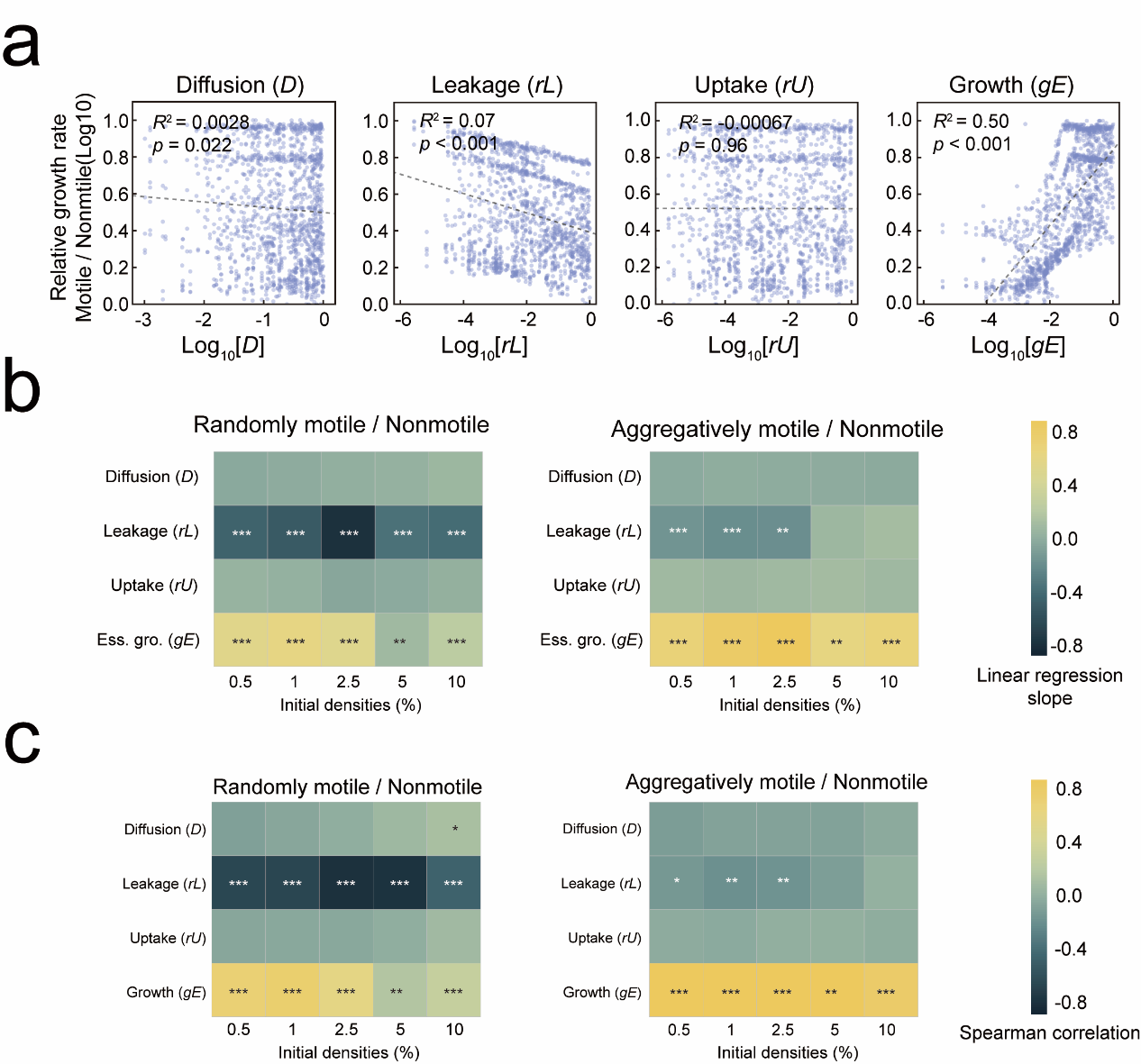

**Figure S9 | Effects of *D*, *rL*, *rU*, and *gE* on the relative population growth rates of motile consortia compared with the nonmotile consortium.**

(a) Linear correlation analyses between *D*, *rL*, *rU*, and *gE* and the relative population growth rates of aggregative motile consortia compared with the nonmotile consortium. (b-c) Effects of *D*, *rL*, *rU*, and *gE* on the relative population growth rates of random motile (left) and aggregative motile (right) consortia compared with the nonmotile consortium under five different initial cell densities, assessed by the slopes from linear regression (b) and the Spearman correlation (c). Statistical significance is indicated as *: *p* < 0.05; **: *p* < 0.01; ***: *p* < 0.001.

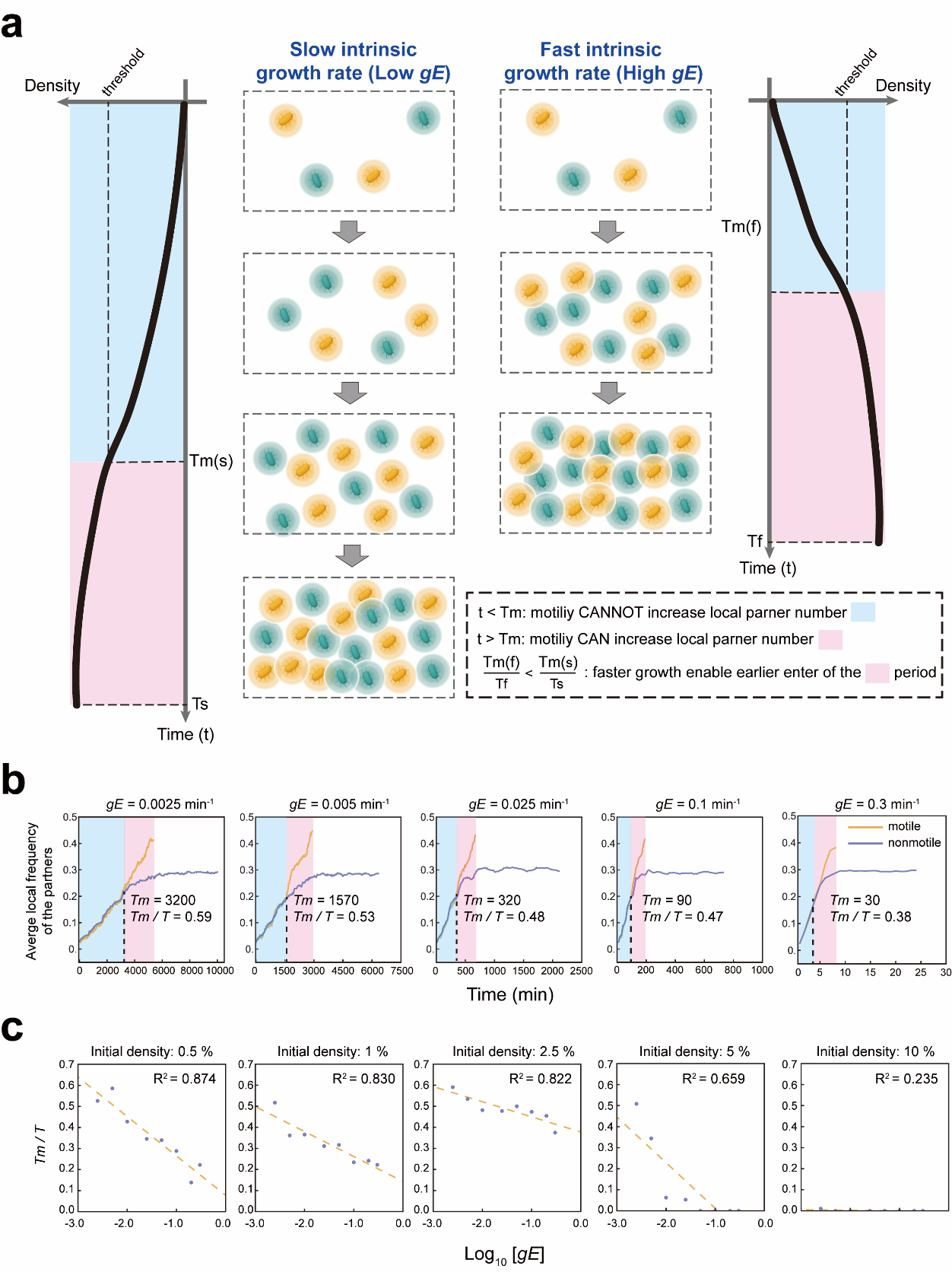

**Figure S10 |** **Effects of intrinsic growth rate on the positive effects of random cell motility.**

(a) A schematic diagram describes how the intrinsic growth rate (*gE*) affects the positive effects of random motility. At the early phase (denoted by the blue region), most cells are surrounded by empty space, so motility rarely leads to encounters with other cells because cell density is low. During this stage, motility has little effect on increasing spatial mixing. As cell density increases over a threshold, after the time point *Tm*, cells begin to encounter nearby neighbours, and motility can then promote the encounter of interacting partners until the motile cells occupy all grids (total time *T*). The pink region denoted this phase. Faster growth shortens the early phase denoted by blue, which can be quantified by a lower value of the ratio *Tm/T*, and thus allows motility to become effective earlier in promoting partner encounters. (b) Temporal dynamics of the average local partner frequency under five different *gE* values. Local frequency was calculated by counting genotype 2 cells within a fixed neighborhood of radius five grids around each genotype 1 cell, dividing this number by 24 (the total neighboring grids), and averaging across all genotype 1 cells. *Tm* was defined as the first time point when the average local partner frequency of the motile consortium exceeded that of the nonmotile consortium. Here, the default parameter values (Table S2) were used for simulations for the random motile and nonmotile consortium. (c) Negative correlation between the *gE* and *Tm*/*T* across five different initial cell densities, where *T* denotes the time used for occupying all grids by the motile consortium. All simulations were performed using the default parameter values listed in Table S2.

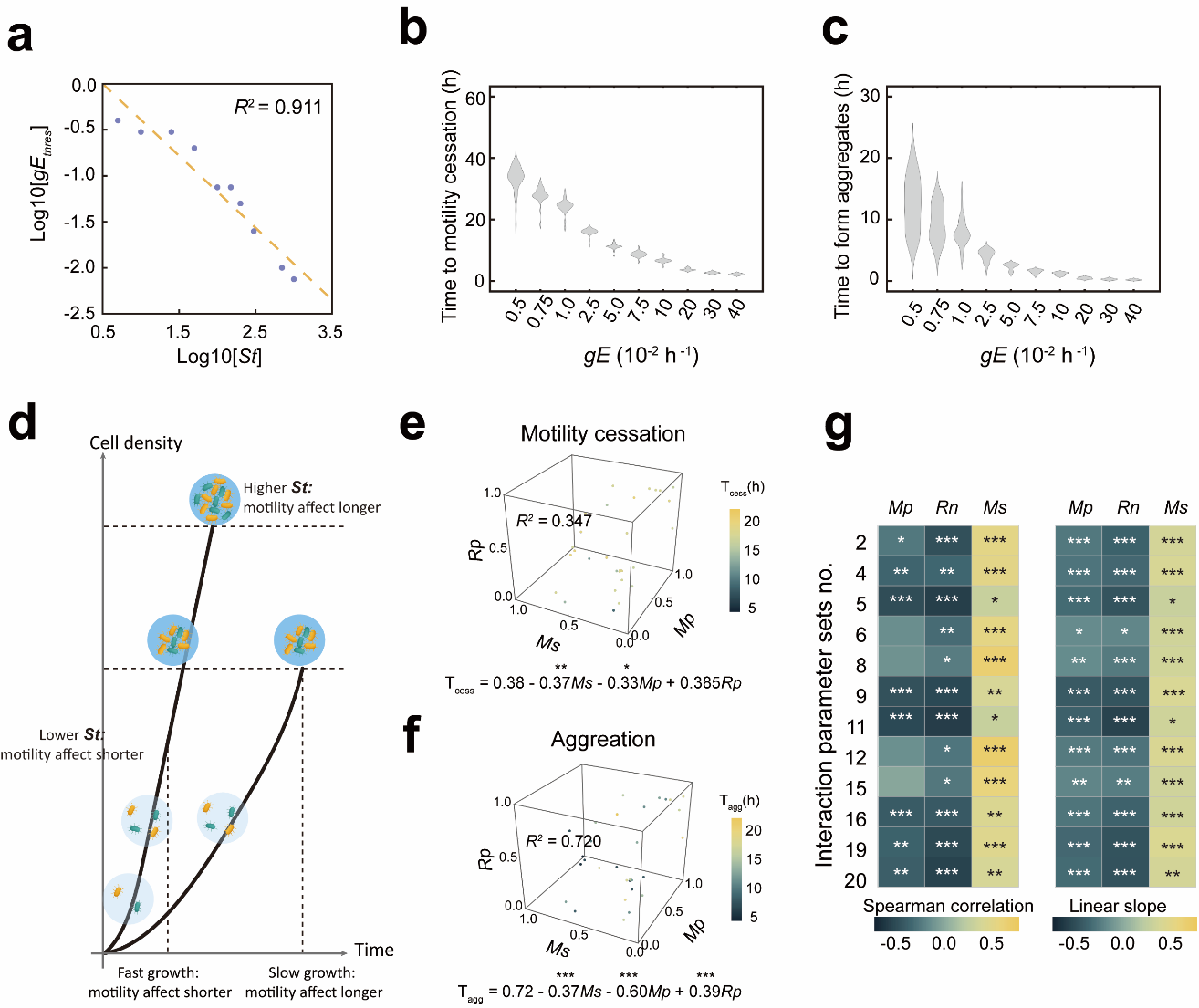

**Figure S11 | Effects of intrinsic growth rate and motility-related parameters on the aggregative behaviours and positive effects of the aggregately motile consortia.**

(a) Negative correlation between the *gE* threshold and the quorum-sensing signal threshold (*St*) that determined motility arrest. (b) Distribution of the first time point at which 90% of cells ceased movement (*T_cess_*) under different intrinsic growth rate (*gE*) values. (c) Distribution of the time points at which cells fully occupied a 40 × 40 grid to form aggregates under different *gE* values. (d) A schematic diagram describes how the intrinsic growth rate (*gE*) and the quorum-sensing signal threshold (*St*) determine the positive effects of aggregative motility. Lower *gE* or higher *St* prolongs the phase during which cells can move and thus strengthens the positive effects of motility on the spatial mixing and community growth. (e–f) Effects of movement probability (*Mp*), reorientation noise (*Rn*), and movement speed (*Ms*) on *T_cess_* (e) and on *T_agg_* (f), the time point at which cells fully occupied a 40 × 40 grid to form aggregates. Results of the multivariate regression analyses are shown below each plot. Statistical significance is indicated above the corresponding parameters as *: *p* < 0.05; **: *p* < 0.01; ***: *p* < 0.001. (g) Effects of *Mp*, *Rn*, and *Ms* on relative growth rates, assessed by Spearman correlation (left) and linear regression slopes (right). Statistical significance: *: *p* < 0.05; **: *p* < 0.01; ***: *p* < 0.001.

### S4 Legends for Supplementary videos

**Supplementary Video 1 |** Time-lapse fluorescence microscopy of the motile consortium (left) and the nonmotile consortium (right) grown in representative 60-μm-diameter circular microfluidic chambers. The △*trpB* cells were labeled with mCherry (shown in yellow pseudocolor), and the △*hisD* cells were labeled with GFP (shown in cyan pseudocolor). Images were captured every 12 minutes.

**Supplementary Video 2 |** Time-lapse fluorescence microscopy of the motile consortium (left) and the nonmotile consortium (right) grown in representative 60-μm square microfluidic chambers. The △*trpB* cells were labeled with mCherry (shown in yellow pseudocolor), and the △*hisD* cells were labeled with GFP (shown in cyan pseudocolor). Images were captured every 12 minutes.

**Supplementary Video 3 |** Time-lapse fluorescence microscopy of the motile consortium (left) and the nonmotile consortium (right) grown in representative 40-μm-diameter circular microfluidic chambers. The △*trpB* cells were labeled with mCherry (shown in yellow pseudocolor), and the △*hisD* cells were labeled with GFP (shown in cyan pseudocolor). Images were captured every 12 minutes.

**Supplementary Video 4 |** Time-lapse fluorescence microscopy of the motile consortium (left) and the nonmotile consortium (right) grown in representative 40-μm square microfluidic chambers. The △*trpB* cells were labeled with mCherry (shown in yellow pseudocolor), and the △*hisD* cells were labeled with GFP (shown in cyan pseudocolor). Images were captured every 12 minutes.

**Supplementary Video 5 |** Time-lapse lineage tracking of the nonmotile consortium grown in microfluidic chambers, showing histidine auxotrophs (left) and tryptophan auxotrophs (right). Cells derived from the same founder are assigned identical colors and number codes based on lineage tracking across divisions. Original time-lapse images are provided in Supplementary Video 1 (right), and corresponding lineage trees are shown in Figure S2b.

**Supplementary Video 6 |** Representative time-lapse sequences showing the motile (top row) and nonmotile (bottom row) consortia. For each row, the left panels display the original fluorescence microscopy images, the middle panels show the corresponding segmented images, and the right panels show quantified single-cell growth rates visualized by color coding, with brighter hues indicating faster growth and the grey color indicating the mistraced cells. The tryptophan auxotroph is shown in yellow and the histidine auxotroph in cyan. Analyses were performed after approximately 65% of the chamber area was occupied (35 frames for the motile consortium and 100 frames for the nonmotile consortium). The full-time-lapse image sequences are provided in Supplementary Video 1.

**Supplementary Video 7 |** Representative simulations showing the spatial and temporal dynamics of the nonmotile (left), randomly motile (middle), and aggregately motile (right) consortia. All simulations were initialized using the default parameter values listed in Table S2.

**Supplementary Video 8 |** Representative simulations showing the dynamics of the aggregately motile consortium under two distinct movement probability (*Mp*) values. The right panel shows the dynamics of the intermixing index. All other parameters were set to the default values listed in Table S2.

**Supplementary Video 9 |** Representative simulations showing the dynamics of the aggregately motile consortium under two distinct movement speed (*Ms*) values. The right panel shows the dynamics of the intermixing index. All other parameters were set to the default values listed in Table S2.

**Supplementary Video 10 |** Representative simulations showing the dynamics of the aggregately motile consortium under two distinct reorientation noise (*Rn*) values. The right panel shows the dynamics of the intermixing index. All other parameters were set to the default values listed in Table S2.
